# An ancestral transmembrane transcription factor couples cell envelope regulation and the SOS response in *Caulobacter crescentus*

**DOI:** 10.64898/2025.12.09.693126

**Authors:** Kamilla Ankær Brejndal, Nikolaj Vestergaard Hansen, Koyel Ghosh, Sebastian Nielsen, Lykke Haastrup Hansen, Lene A. Jakobsen, Martin R. Larsen, Clare L. Kirkpatrick

## Abstract

The DNA damage (SOS) response in bacteria involves derepression of a set of genes in order to activate mechanisms to tolerate stress, repair DNA and slow down cell division. While some of these genes are well characterized, many genes exist which are clearly induced by DNA damage but for which the function is unclear. In *Caulobacter crescentus*, the toxin-antitoxin (TA) system *higBA* and a closely associated downstream transcription factor (*higX*) are strongly induced as part of the SOS response, but the role of *higX* is unknown. We show that, unexpectedly, HigX functions independently of HigBA as a cell membrane-associated regulator and is toxic when overexpressed in filamentous cells. ChIP-Seq indicated that it binds to several promoters associated with cell envelope regulation. In cells with the SOS response constitutively activated, HigX was overproduced, but at the same time was unable to bind the majority of its target promoters. *higX* was highly conserved in genomic context among many alpha-proteobacteria, while *higBA* was only found upstream of it in a small number of *Caulobacter* genomes, including the universal laboratory strain *C. crescentus* NA1000. Compositional analysis suggested that *higBA* originated from a foreign source, while *higX* is likely ancestral to the alpha-proteobacteria. Our data support a model where dysregulation of HigX production and activity in filamentous Δ*lexA* cells contributes to cell envelope instability and antibiotic sensitivity. Thus, the protective effect of inhibiting cell division during the SOS response in order to repair the DNA, carries the hidden cost of interference with HigX-mediated cell envelope maintenance.

**Author summary:** The DNA damage (SOS) response in bacteria is key for their survival in harsh conditions, but the genes which are induced by this response and the molecular mechanisms involved are variable between different bacteria. We investigated the function of an uncharacterized SOS-induced gene in *Caulobacter crescentus* (*higX*) and found that it is a membrane-localized DNA binding protein, but that it did not help survival in DNA damage. Instead, it negatively affected cell viability when induced, unlike other SOS response genes. Genomic analysis showed that *higX* is highly conserved among alphaproteobacteria, but it is only under SOS control in a small number of genomes (including *Caulobacter crescentus*) where a toxin-antitoxin system and its SOS-regulatory promoter have been inserted upstream. This work uncovers HigX as a novel factor in cell envelope regulation, and shows that loss of cell shape regulation, eg. in filamentous Δ*lexA* cells, can influence the ability of transmembrane transcription factors to bind to their target genes.

## Introduction

Free-living bacteria require robust stress response systems in order to thrive in variable (and sometimes stressful) natural environments. One of the most important is the SOS response system, for response to and repair of DNA damage, especially double-strand breaks (1). When DNA damage occurs, the SOS response allows cell division to be paused for the cell to carry out DNA repair, in order to assure that intact chromosomes are passed on to the next generation. LexA and RecA are the key regulators of the SOS response, and are widely conserved among bacteria (2). In a noninduced state, LexA maintains its target genes under tight repression by binding to specific recognition sites in their promoters. During DNA damage, RecA recognizes single-stranded DNA and polymerises into a filament (3, 4). The polymer form of RecA interacts with LexA causing it to cleave itself and leave the promoters, allowing transcription of the target genes (5). While the consensus binding sequence for LexA varies between different bacterial species, it is generally true that the closer a given LexA binding site matches the consensus, the higher the affinity of LexA for the site and the tighter the repression of the promoter(6). Hence, a high-affinity site can act as an on-off switch for its target genes, while a low-affinity site would have a more subtle effect on transcription.

Toxin-antitoxin (TA) systems are widespread in bacterial genomes, with a tendency towards free-living bacteria possessing multiple systems (frequently in multiple copies) and intracellular bacteria having few or none (7). All systems consist of at least a toxin, which acts intracellularly to inhibit central metabolism, and an antitoxin which inhibits toxin action (8–11). In some cases, described further later, a third component is also present. The regulation of TA system activity is primarily post-translational, because while the promoters of some TA systems are transcriptionally activated by stress (12–15), transcriptional activation does not automatically result in a net increase of toxin activity (16) and other systems are transcriptionally stress-insensitive (17, 18). In some bacteria, TA system promoters are under the control of LexA (12–15)￼. One example is the *higBA* system of *Caulobacter crescentus*. This TA system comprises the DNA-binding protein antitoxin HigA and the ribosome-dependent RNase toxin HigB and is negatively regulated by binding of HigA and LexA upstream of the *higB* start codon. LexA binding to the *higBA* promoter provides sufficiently strong repression that it is possible to delete the HigA antitoxin without major phenotypic effects(15)￼. Consistent with this, the LexA binding site in the *higBA* promoter is a perfect match to the experimentally defined consensus sequence for *Caulobacter* LexA (19)￼ (T/A)TGTTC (N_6_)TGTTC, and the HigA protein is almost undetectable in wild type (WT) *Caulobacter crescentus* but strongly upregulated in the Δ*lexA* mutant (15)￼. Due to this strict repression, HigB-dependent phenotypes are most clearly observable in the Δ*lexA* mutant. In this strain, improvements in growth rate and in ciprofloxacin resistance are seen when the *higB* toxin gene is deleted, suggesting that the *higBA* TA system is a functional component of the *Caulobacter crescentus* DNA damage response and that HigB’s toxic activity contributes to poor growth rate and ciprofloxacin sensitivity in Δ*lexA* cells. Hence, *Caulobacter crescentus higBA* is an example of a TA system in which transcriptional derepression of its promoter (during DNA damage) does lead to toxin-dependent phenotypes(15)￼.

One toxin-dependent phenotype of the HigBA system was an unexpected protective effect against cell envelope stress. This came to light through a forward genetic screen for rescue of nalidixic acid sensitivity in the Δ*tipN* mutant, which identified *higA* as a loss of function suppressor candidate, during a transposon insertion screen for nalidixic acid resistance (15). The mechanism of this protective effect proved to be through HigB-mediated cleavage of the *acrB2* efflux pump transcript mRNA, which was strongly transcriptionally induced by nalidixic acid. Mutants lacking TipN had destabilized cell envelopes which were disproportionally sensitive to cell envelope stress induced by overload of the envelope with AcrAB2 efflux pump proteins (20), unless these were post-transcriptionally downregulated by HigB.

While our previous work focused on the role of HigB in the DNA damage and cell envelope stress responses, it also came to our attention that other factors may be involved. In particular, we noted different antibiotic resistance phenotypes for the Δ*lexA* Δ*higB* and the Δ*lexA* Δ*higBA* mutants, even though both of these are lacking the toxin gene *higB*. The Δ*lexA* Δ*higBA* strain remained as sensitive to DNA damaging antibiotics as the Δ*lexA* parent strain, and did not display the improved resistance seen in the Δ*lexA* Δ*higB* mutant (15). To investigate sources of this HigB-independent antibiotic sensitivity, we examined the genomic context of the *higBA* TA system and observed that a third gene is encoded only 42 bp downstream of the *higA* antitoxin gene. The close proximity of this gene (CCNA_03131) to *higBA* was suggestive of a function as a regulatory third component of the TA system. However, it bore no similarity to the chaperone third components seen in other *higBA* systems, in some cases designated *higC* (21, 22). Instead, it is annotated as a transcription factor of the LytTR family. These factors are found in a wide variety of bacteria, including pathogens, and regulate processes such as competence, extracellular toxin and bacteriocin production, alginate production and quorum sensing (23). Here we show that this gene, for which we propose the designation *higX*, (considering its close association with *higBA* but still enigmatic function) is indeed a transcription factor and can regulate the TA system promoter, but its primary function is independent from it. In evolutionary terms, *higBA* in *Caulobacter crescentus* is a recent arrival upstream of *higX*. Insertion of *higBA* and its LexA-dependent promoter at this position is unique to the *crescentus/vibrioides* lineage and is not conserved among other *Caulobacter* species or other alpha-proteobacteria. Conversely, *higX* is widely conserved among the alphaproteobacteria, generally found adjacent to a putative hydrolase gene and is not LexA-dependent. Overproduction of *higX* upon LexA derepression in *Caulobacter crescentus* leads to ciprofloxacin sensitivity, likely because the cell envelope is destabilized and the antibiotic has easier access to its target inside the cell. However, we also observed that in the Δ*lexA* strain, binding of HigX to its target promoters was globally decreased, suggesting a novel link between cell filamentation and cell envelope gene regulation.

## Results

### Ciprofloxacin sensitivity of the ΔlexA ΔhigBA mutant is due to excessive production of a transcription factor downstream of higBA

To investigate the source of the unexpected HigB-independent ciprofloxacin sensitivity in Δ*lexA* Δ*higBA*, we focused on the CCNA_03131 (hereafter, *higX*) gene downstream of *higBA* (Fig. 1A). We constructed in-frame deletions of *higX* in WT, Δ*lexA* or Δ*higBA* strains and investigated the effect on ciprofloxacin sensitivity. Consistent with our previous observations (15), loss of *higA* in the WT background sensitized cells to ciprofloxacin (through HigB activation), while deletion of *higB* from the Δ*lexA* strain was protective against it. Deletion of *higX* in WT, with or without deletion of *higBA*, had no effect. However, in the ciprofloxacin-sensitive Δ*lexA* Δ*higBA* background, deletion of *higX* improved ciprofloxacin resistance to a similar level to the Δ*lexA* Δ*higB* strain, showing that *higX* is indeed responsible for the ciprofloxacin sensitivity in the Δ*lexA* Δ*higBA* strain in the absence of the HigB toxin (Fig. 1B).

**Figure 1.**
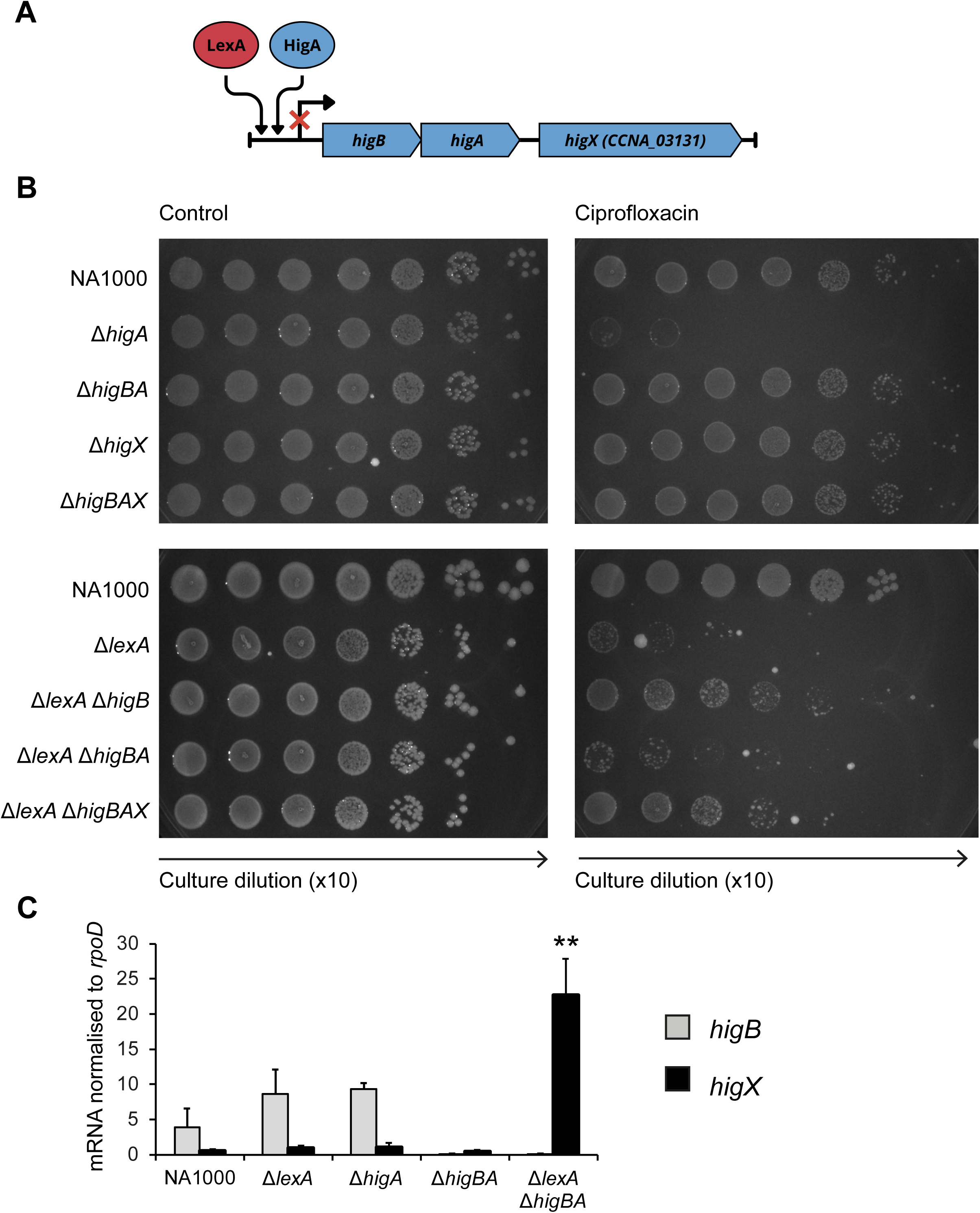
The higBA toxin-antitoxin system has a transcription factor closely associated with it. (A) Operon structure of *higBA* and location of the downstream gene *CCNA_03131* (*higX*). The promoter upstream of *higBA* is negatively regulated by LexA and HigA. The *higB* and *higA* genes are translationally coupled, with 7 bp overlap between the stop codon of *higB* and the start codon of *higA*, while the *higX* start codon is 42 bp downstream of the *higA* stop codon. (B) Efficiency of plating assay of WT *Caulobacter* (NA1000) and mutant strains lacking combinations of *lexA*, *higB*, *higA* and/or *higX*, on 0.5 μg/ml ciprofloxacin or vehicle control. Images are representative of three independent biological replicates. (C) Quantitative RT-PCR of mRNA levels of *higB* and *higX*, normalized to the control transcript *rpoD*. Data was obtained from three independent biological replicates.

The Δ*lexA* Δ*higBA* strain had been constructed by in frame deletion of *higBA* in the Δ*lexA* single mutant background. However, we subsequently realized that this would exert polar regulatory effects on *higX* expression, because (1) deletion of the *higBA* coding sequence brings *higX* directly downstream of the strong promoter regulating *higBA*, and (2) both repressors of this promoter, LexA and HigA, are absent. Indeed, we observed that *higX* was transcribed at very low levels in WT, Δ*higA* and Δ*higBA* but much higher in Δ*lexA* Δ*higBA* (Fig. 1C). Therefore, the ciprofloxacin sensitivity phenotype of Δ*lexA* Δ*higBA* is associated with elevated HigX RNA expression, suggesting why *higX* deletion is protective.

### HigX regulates the expression of higBA alongside LexA and HigA

In other three-component TA systems with a transcription factor as their third component, the transcription factor regulates of the promoter upstream of the TA system (18, 24). To investigate whether HigX also has this activity, we examined activity of a transcriptional P*_higBA_-lacZ* reporter construct (Fig. 2A, B) in WT or strains lacking combinations of *lexA*, *higBA* or *higX*. Promoter activity was unaffected by *higX* deletion in the WT or in Δ*higBA* backgrounds, but decreased in Δ*lexA* Δ*higBAX* compared to Δ*lexA* Δ*higBA*, suggesting that HigX can activate the promoter upstream of *higBA* when LexA and HigA are absent (Fig. 2B). We attempted to confirm this finding by complementation with an overexpression construct containing *higX* under the control of the vanillate-inducible promoter (pMT335-*higX*). However, we could not transform the Δ*lexA* Δ*higBA* strain with this plasmid, suggesting that its increased level of *higX* expression is sufficiently toxic to make the cells intolerant of transformation with a multi-copy plasmid containing *higX*, even in the absence of inducer. Expression of *higX* from this plasmid in WT, Δ*lexA* or Δ*higBA* strains containing the P*_higBA_-lacZ* transcriptional reporter had a weak inhibitory effect on promoter activity, which was only statistically significant in the Δ*higBA* background (Fig. 2C). Therefore, *higX* can activate or repress the promoter upstream of *higBA*, dependent on which other regulators are present, but its regulatory activity is weaker at this locus than that of LexA or HigA.

**Figure 2.**
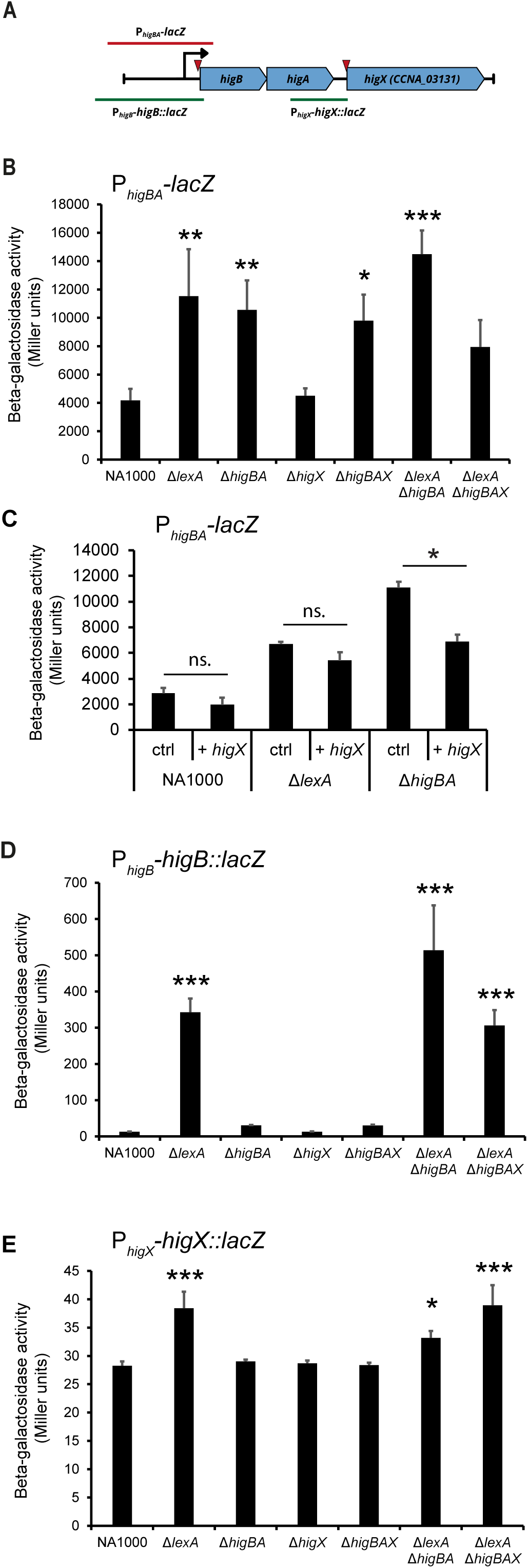
Translational reporter constructs show that higB, higA and higX are independently regulated. (A) Construction of the transcriptional (red) or translational (green) reporter fusions to lacZ from regions of the *higBAX* genomic locus. The transcription start sites in this region previously detected by global mapping (25) are shown as red arrows. (B-C) Transcriptional beta-galactosidase reporter assays in WT and mutant strains lacking combinations of *lexA*, *higB*, *higA* and/or *higX* (B) and WT, Δ*lexA* and Δ*higBA* strains containing the empty vector pMT335 or the *higX* overexpression plasmid pMT335-*higX*. Overexpression of *higX* was induced by 50 μM vanillate (C). (D-E) Translational beta-galactosidase reporter assays in WT and mutant strains lacking combinations of *lexA*, *higB*, *higA* and/or *higX* for a *lacZ* in-frame fusion to the *higB* upstream region and first ten codons of HigB (D), and a *lacZ* in-frame fusion to the *higX* upstream region and first four codons of HigX (E). A promoterless negative control of the translational fusion construct was tested in WT and had undetectable beta-galactosidase activity (zero to two Miller units).

The different levels of mRNA expression for *higB* and *higX* observed by qRT-PCR (Fig. 1C) were suggestive of regulation of the two transcripts by separate promoters, consistent with previous genome-wide transcriptional profiling which identified independent primary transcriptional start sites for *higBA* and *higX* (25), at genome positions 3281528 and 3280865 respectively. We therefore constructed translational beta-galactosidase fusions of the regions immediately upstream of *higB* and *higX* (including the first few codons of each of the genes) to *lacZ*, such that the *higB* and *higX* reporter fusions should employ the native transcription start sites,. Surprisingly, the translational fusion P*_higB_-higB::lacZ* did not exhibit the same pattern of activity as the transcriptional reporter (Fig. 2D, compare to 2B). For the translational reporter, the basal activity in WT, Δ*higBA*, Δ*higBAX* or Δ*higX* strains was almost undetectable and high promoter activity was only observed in the three strains lacking *lexA*, in contrast to the transcriptional reporter where basal expression appeared high and the fold increase in the Δ*lexA* mutant strains was much less pronounced. Conversely, the activity of the translational reporter for *higX* expression was much weaker (Fig 2E). It was slightly LexA-dependent, but much less so than the promoter of *higB.* We conclude that *higX* is primarily regulated by the strictly LexA-dependent promoter upstream of *higB*, despite having an independent transcription start site, and that the different mRNA levels observed for *higB* vs. *higX* (Fig. 1C) are due to other factors, for example processing or different stability. Moreover, the strikingly low basal level of P*_higB_-higB::lacZ* activity, relative to the transcriptional *lacZ* fusion of the same region, is suggestive of an unknown post-transcriptional negative regulator of *higBA*.

### HigX is a membrane-associated transcription factor

It was previously noted in a bioinformatic study of LytTR-family transcription factors (23) that all members of this family in the *Caulobacter crescentus* genome, including *higX*, are likely to encode transmembrane proteins. Bioinformatic prediction suggested that HigX contains four transmembrane helices N-terminal to the LytTR DNA binding domain, with the transmembrane and DNA-binding domains separated by a short low-complexity region (LCR) (Fig. 3A, S1A). The four helices are predicted to form a tightly packed bundle with no apparent periplasmic domain (Fig. S1B). To investigate whether the transmembrane domain was required for DNA binding activity, we constructed another overexpression plasmid where the *higX* gene was truncated to convert amino acid 133 (Val) to a new start codon, creating the *higX*-noTM variant containing only the LCR and the LytTR domain. We expressed this alongside the full-length *higX* overexpression plasmid and compared HigX abundance by western blotting using a custom polyclonal antibody to HigX alongside cell extracts containing pMT335 empty vector and NA1000 without any plasmid as control. It was only possible to detect HigX protein in the pMT335-*higX* overexpression strain, showing that while the antibody can detect HigX, the native level of this protein is very low (Fig. 3B, left). We could not detect the truncated HigX protein from the pMT335-*higX*-noTM overexpression strain, suggesting that it may be unstable. Comparing the *Caulobacter* western blot results to cell extracts of *E. coli* Rosetta cells containing pET28a-*higX* or pET28a-*higX*-noTM constructs intended for protein purification, we observed that both the full-length and truncated proteins could be detected in *E. coli* with our HigX polyclonal antibody (Fig. 3B, right). Therefore, although the truncated version of HigX is not intrinsically unstable, it cannot be maintained in *Caulobacter* in the absence of the TM domain.

**Figure 3.**
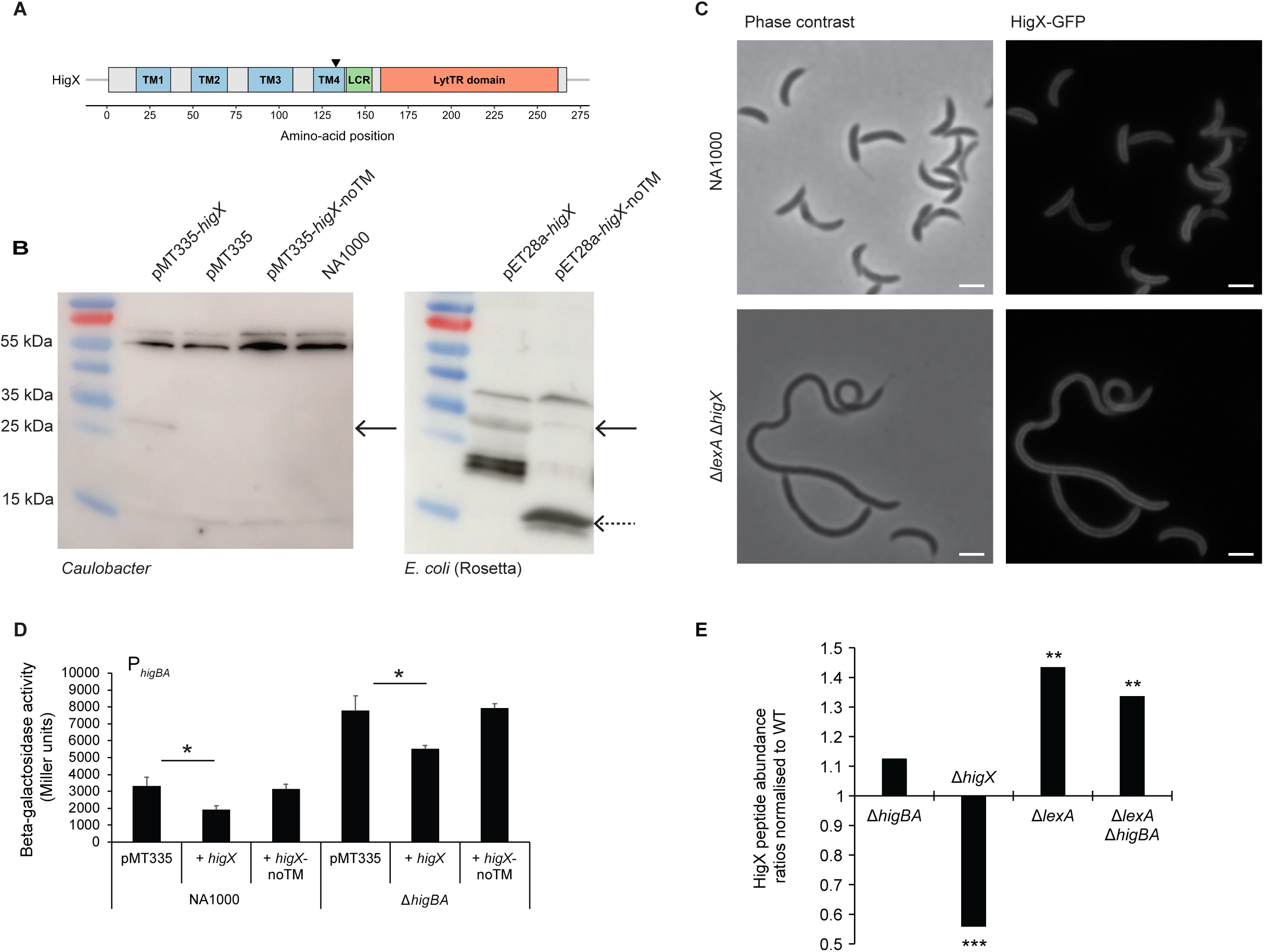
The N-terminal region of HigX localizes it to the cell periphery and is required for higBA promoter regulatory activity. (A) Predicted domain structure (from Uniprot) of HigX with transmembrane helices (TM1-4) indicated in blue, low-complexity region (LCR) in green and the LytTR DNA-binding domain in orange. The black pin indicates where in the *higX*-noTM construct, amino acid Val133 was converted to a new Met start codon to create a truncated version consisting of the DNA binding domain without any transmembrane helices. (B) Western blots of cell extracts of *Caulobacter* cells expressing *higX* or *higX*-noTM in pMT335 with 50 µM vanillate induction (left) or E. coli Rosetta expressing His6-HigX in pET28a with 0.1 mM IPTG induction (right), with the anti-HigX polyclonal antibody. Solid arrows indicate the size of full-length HigX (28.2 kDa) and the dotted arow indicates the size of HigX-noTM (14.6 kDa). (C) Phase-contrast and fluorescence microscopy of WT and Δ*lexA* Δ*higX* expressing a HigX-GFP translational fusion from the vanillate-inducible promoter. Scale bars indicate 2 μm. No GFP fluorescence was observed for cells carrying the pMT335 empty vector. (D) Beta-galactosidase assay of the transcriptional P*_higBAX_*-lacZ reporter in WT and Δ*higBA* strains containing the empty vector pMT335, the full length *higX* overexpression plasmid pMT335-*higX* or the truncated *higX* overexpression plasmid pMT335-*higX*-noTM. No statistically significant difference was observed between the empty vector control and the *higX*-noTM overexpression plasmid. (E) Relative abundances of HigX peptides in Δ*higBA*, Δ*higX*, Δ*lexA* and Δ*lexA* Δ*higBA* membrane-enriched cell extracts detected by mass spectrometry and normalized to WT. Values are the averages of three independent biological replicates. Peptide abundance values (averages), CV% values (% coefficient of variation) and p values are in Supplementary Table S1. Note that non-zero values for HigX peptides in the Δ*higX* strain are likely derived from the vestiges of the start and end of the *higX* coding region that remain in place after the in-frame deletion, and/or low-frequency detection of peptides derived from other proteins that coincidentally have the same mass/charge ratio as HigX-derived peptides.

To investigate whether HigX is localized to the membrane in living cells, we constructed C-terminal GFP fusions to full length HigX and to the truncated HigX-noTM allele lacking the transmembrane helices. Since the native level of HigX expression was so low, these were created as plasmid-borne constructs under control of the vanillate-inducible promoter in order to have sufficient expression to detect GFP fluorescence. Fluorescence microscopy showed that full length HigX localizes uniformly around the cell periphery in both WT and filamentous cells (Fig. 3C), indicating that normal cell division is not required for HigX subcellular localisation. No fluorescence was observed in cells carrying the truncated HigX-noTM-GFP fusion protein, and overexpression of the truncated HigX-noTM protein (without GFP fusion) had no effect on activity of the P*_higBA_-lacZ* transcriptional reporter (Fig. 3D), confirming that the transmembrane domain is essential for stability and/or activity of HigX in the cell.

In order to detect the HigX native expression levels in WT and mutant strains, we employed quantitative mass spectrometry. Protein extracts were prepared from WT, Δ*higBA*, Δ*higX*, Δ*lexA*, and Δ*lexA* Δ*higBA* strains, with separation of the soluble (cytoplasmic) proteins from the insoluble (membrane-associated) proteins before trypsinization and isotope labeling. A total of 1332 membrane-associated proteins were detected with two or more peptides, including HigX. No HigX peptides were detected in the soluble fraction, indicating that the fluorescence observed at the cell periphery in Fig 3C is due to membrane localisation. The abundance of HigX peptides was not significantly changed between WT and Δ*higBA*, strongly depleted in Δ*higX* and enriched in both Δ*lexA* and Δ*lexA* Δ*higBA* strains by 30-40% (Fig. 3E, Supplementary Table S1), consistent with the trend observed in the translational reporter experiments (Fig. 2).

### HigX acts independently of the toxin-antitoxin system HigBA

To test our hypothesis that overproduction of HigX was responsible for ciprofloxacin sensitivity in Δ*lexA*, we expressed it *in tran*s from the vanillate-inducible promoter using pMT335-*higX* in WT, Δ*lexA* and Δ*lexA* Δ*higB*. We found that *higX* overexpression had no effect on cell viability compared to the empty vector control on normal media. However, on ciprofloxacin, *higX* overexpression reduced viability of the Δ*lexA* strain compared to the empty vector. Strikingly, the *higX* overexpression plasmid also reversed the improvement in ciprofloxacin resistance seen in the Δ*lexA* Δ*higB* strain relative to empty vector (Fig. 4A). In addition to confirming that *higX* overexpression was responsible for the ciprofloxacin sensitivity phenotype, this result also indicated that HigX does not act solely through regulation of the TA system promoter, since the phenotype was clearly visible in the absence of the HigB toxin. In support of the independence of HigX from HigBA, we also observed an improvement of ciprofloxacin resistance in a Δ*lexA* Δ*higX* double mutant relative to ΔlexA, where both of these strains have the *higBA* TA system genes intact (Fig. 4B). This result demonstrates that the increased abundance of HigX seen in the Δ*lexA* single mutant strain (Fig. 3D) contributes to its ciprofloxacin sensitivity in addition to (and independently of) the overexpression of HigB.

**Figure 4.**
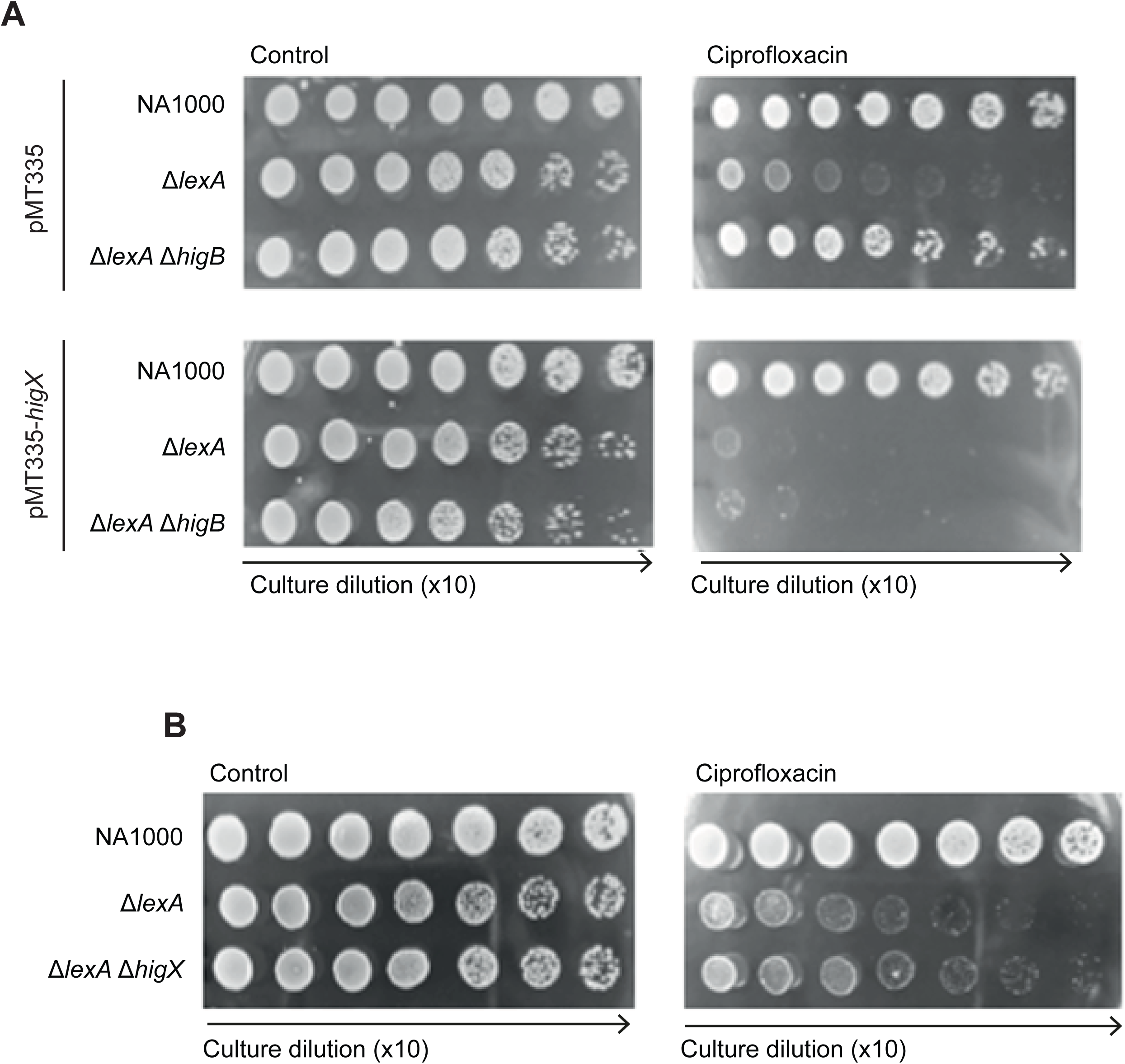
Overexpression of higX in trans causes ciprofloxacin sensitivity independently of the higBA toxin-antitoxin system. (A) Efficiency of plating assay of WT, Δ*lexA* and Δ*lexA* Δ*higB* containing empty vector pMT335 or the *higX* overexpression plasmid pMT335-*higX*. All plates contained gentamicin and 50 μM vanillate in addition to ciprofloxacin (0.5 μg/ml) or vehicle control. (B) Efficiency of plating assay of WT, Δ*lexA* and Δ*lexA* Δ*higX* on ciprofloxacin (0.5 μg/ml) or vehicle control. Images are representative of three independent biological replicates.

### HigX preferentially binds cell envelope, transport, and metabolism-associated genes

To assess the DNA binding capability of HigX, we performed chromatin immunoprecipitation sequencing (ChIP-Seq) on wild type and Δ*lexA* strains using our polyclonal antibody to HigX. Using the “MACS2 callpeak” program, we identified 596 significantly enriched peaks in the wild-type strain (Supplementary Dataset S1). In contrast, peak calling for the Δ*lexA* strain revealed genome-wide depletion of the enriched peaks, with 413 peaks identified with significant enrichment, but with enrichment signal values uniformly lower than in WT (Fig 5A). Therefore, the DNA-binding and thus regulatory function of HigX is strongly reduced upon deletion of *lexA*, despite the 40% upregulation of HigX peptide abundance (Fig 3E). Significant peaks in the WT strain were further filtered through a 95% threshold based on their signal value (≥ 3.22), resulting in 30 significant peaks that corresponded to the top 5% of the dataset. The highest of these peaks was located between CCNA_03217 (a TetR-family transcriptional regulator) and CCNA_03218 (periplasmic multidrug efflux lipoprotein precursor), with the summit being closest to CCNA_03218 (Fig 5B).

**Figure 5.**
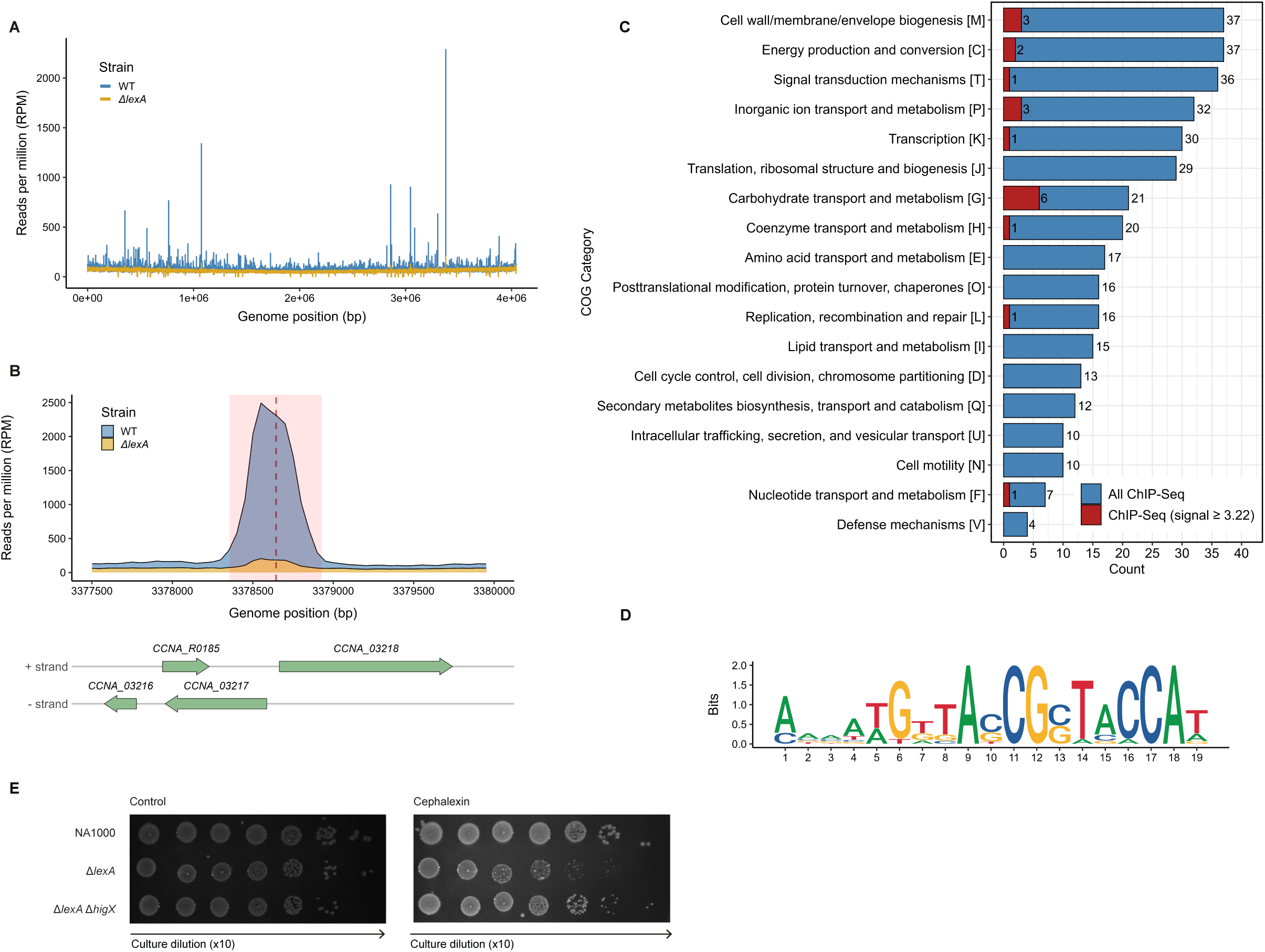
Genome-wide analysis of HigX DNA-binding and functional enrichment. (A) ChIP-seq profiles of HigX in *Caulobacter* NA1000 wild-type (WT, blue) and *ΔlexA* (yellow) strains, with the data plotted as reads per million (RPM) across the genome. (B) Zoom-in of the highest peak in the dataset located between *CCNA_03217* and *CCNA_03218*, with a gene track displaying the transcriptional orientation. The peak region is highlighted in red, and the summit position is indicated by a red dashed line. (C) COG analysis of significant ChIP-seq peaks in the WT strain. Blue bars represent all significant peaks identified by MACS2, while red bars indicate peaks filtered by a signal value threshold (≥ 3.22). Two categories were omitted: “Function unknown (S)” with 66 (blue) and 3 (red) counts, and “Unassigned (no COG)” with 177 (blue) and 9 (red) counts. (D) Sequence logo of the HigX binding motif identified through MEME-ChIP from the peak summit regions of the filtered peaks from the WT (signal value ≥ 3.22). The motif consists of an imperfect inverted repeat of 8-bp with no intervening spacer. The left half of repeat is located at positions 4–11 (ATGTTASC) and the right half at positions 12–19 (GSTACCAT) with a single mismatch between the repeat halves. (E) Efficiency of plating assay of WT, Δ*lexA* and Δ*lexA* Δ*higX* on cephalexin (5 μg/ml) or vehicle control. Images are representative of three independent biological replicates.

To determine the overall function of HigX target genes and provide a holistic view of its potential regulatory role, we extracted the summit locations of all 596 peaks and mapped them to the nearest genes, followed by COG (Clusters of Orthologous Groups) annotation and functional classification. From this, we observed that most peaks were associated with the following COG categories, listed from most to least abundant: M, C, T, P, K, and J (Fig. 5C). However, when we focused on the top 30 significant peaks (signal value ≥ 3.22), genes associated with carbohydrate transport and metabolism (G) became the most prominent, while most of the previously dominant categories (M, C, T, P, and K) remained to a lesser extent. In contrast, genes associated with translation, ribosomal structure, and biogenesis (J) were no longer represented. This COG distribution suggests that, for the 5% highest confidence ChIP peaks, HigX preferentially targets genes associated with membrane, transport, and metabolism-related functions.

To further characterize the DNA-binding specificity of HigX, we performed a motif discovery analysis by utilizing the summits of the 30 significant (top 5%) ChIP-seq peaks in conjunction with MEME-ChIP. This analysis identified a significantly enriched 19-bp motif (E-value = 1.4 × 10⁻^23^) present in 13 of the 30 submitted sequences, with a strong central enrichment, which indicates that this motif likely represents the direct binding site of HigX (Fig. 5D). The motif is characterized by an 8-bp imperfect inverted repeat, with the left half spanning positions 4–11 (ATGTTASC) and the right half spanning positions 12–19 (GSTACCAT), with no intervening spacer and a single mismatch between the repeat halves.

Thus far, we had only investigated ciprofloxacin sensitivity of the Δ*lexA* strain and derivatives. However, the ChIP-Seq analysis suggested that the dysregulation of HigX in the Δ*lexA* strain, either at the level of overproduction of the protein or failure to bind the target genes, may also lead to cell envelope irregularity that could lead to sensitivity to cell envelope-targeting antibiotics. We therefore tested the ability of the Δ*lexA* and Δ*lexA* Δ*higX* strains to grow on cephalexin, which targets septal peptidoglycan synthesis. The Δ*lexA* strain was sensitized to cephalexin relative to WT, and cephalexin resistance was restored in the Δ*lexA* Δ*higX* double mutant (Fig. 5E). Therefore, even though HigX production is under SOS control, its downstream effects in the cell are more likely to be involved with cell envelope maintenance than with the SOS response.

### Suppressor mutations in cell envelope-related genes improve ΔlexA ΔhigBA ciprofloxacin resistance

To identify possible downstream effectors of HigX responsible for the overexpression phenotype, we designed parallel forward genetic screening strategies to take advantage of (1) the Δ*lexA* Δ*higBA* strain’s sensitivity to ciprofloxacin and (2) its intolerance of transformation with the pMT335-*higX* plasmid (Fig 6A). We constructed a pooled library of random *himar1* transposon (Tn) mutants in the Δ*lexA* Δ*higBA* background in order to select for (1) ciprofloxacin-resistant Tn mutants and (2) Tn mutants which can be stably transformed with pMT335-*higX*, using the Δ*lexA* Δ*higBA* strain in preference to the Δ*lexA* strain in order to avoid selection of phenotypes due to the HigB toxin. The first screen was performed by passage of the mutant pool through medium containing 1 μg/ml ciprofloxacin, followed by selection of ciprofloxacin-resistant single colonies. The second was performed by preparation of electrocompetent cells from a culture of the pooled library, electroporation with pMT335-higX and plating on medium containing gentamicin and vanillate (to select for the plasmid antibiotic resistance marker and to induce higX overexpression, respectively). Such mutants should have Tn insertions in loci which suppress the HigX-mediated phenotypes.

**Figure 6.**
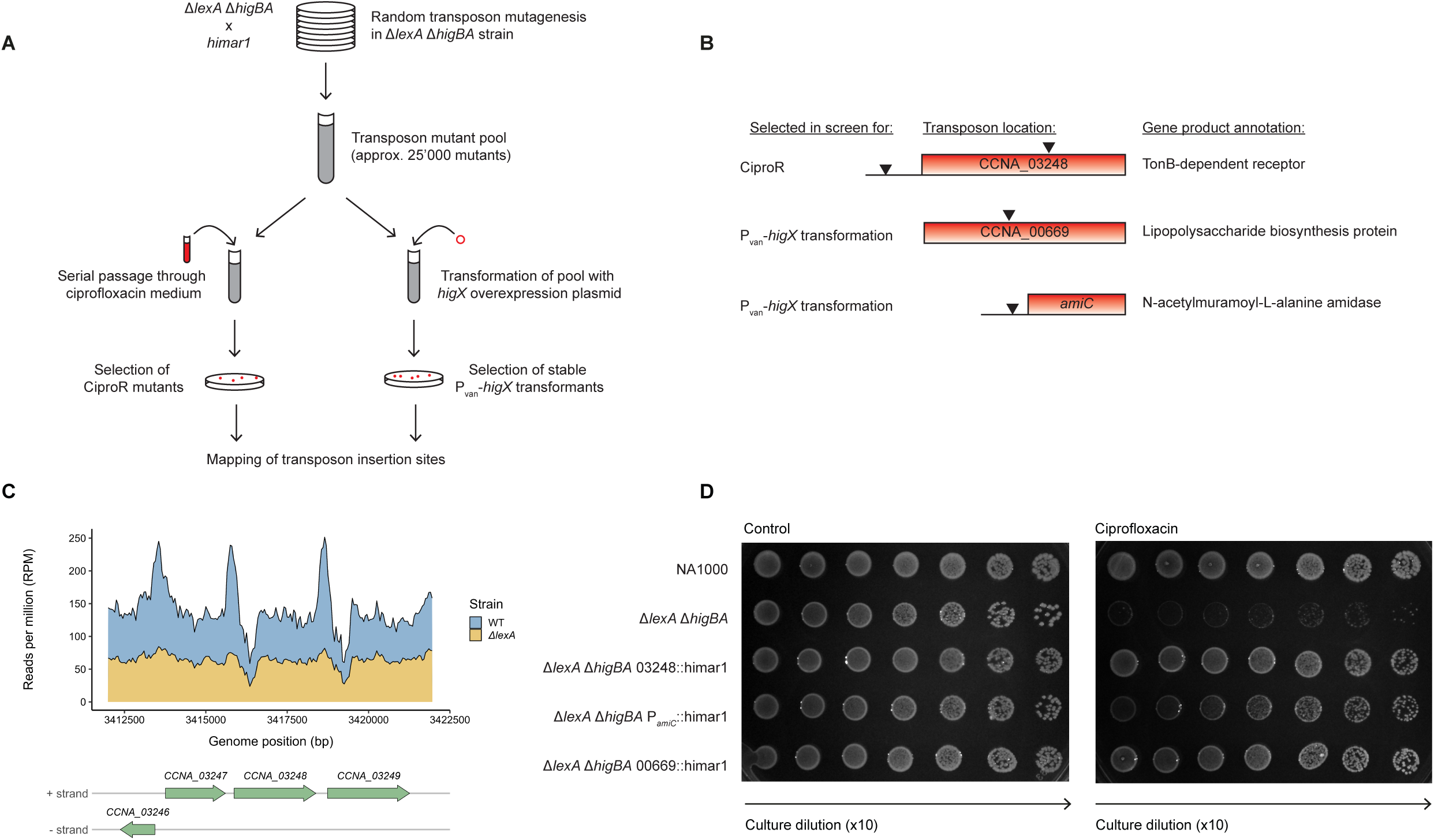
Forward genetic screening for suppressors of ΔlexA ΔhigBA ciprofloxacin sensitivity reveals cell envelope-related genes. (A) Schematic of the forward genetic screening protocol in the Δ*lexA* Δ*higBA* strain for ciprofloxacin resistance (left) and pMT335-*higX* transformation tolerance (right). (B) Location of the transposon insertions in the four verified suppressor mutants. The insertion sites at the CCNA_03248 locus are at 3415650 (upstream) and 3417411 (inside CDS), the insertion site in the CCNA_00669 gene is at 727111 and the insertion site upstream of the *amiC* gene is at 2097088. (C) Zoom-in of the ChIP-Seq profiles of WT (blue) and Δ*lexA* (yellow) at the CCNA_03248 locus. (D) Efficiency of plating assay of WT, Δ*lexA* Δ*higBA* and Δ*lexA* Δ*higBA* transduced with three of the suppressor transposon mutations, on ciprofloxacin (0.5 μg/ml) or vehicle control. Images are representative of three independent biological replicates.

From our Tn mutant library of approximately 25’000 clones, we were able to obtain four stable Tn mutants where we could confirm the desired phenotypes by re-plating on selective media (ciprofloxacin, or gentamicin + vanillate) followed by backcrossing of the transposon mutations into Δ*lexA* Δ*higBA* cells which had not been through the mutagenesis or selection procedure. All four suppressor mutations were in or near genes associated with cell envelope-related functions (Fig. 6B). The two Tn mutants obtained from the ciprofloxacin resistance screen both had a Tn insertion in or near CCNA_03248, encoding an uncharacterized TonB-dependent receptor. One insertion was found within the coding sequence of CCNA_03248, while the other was in an intergenic region 218 bp upstream of the CCNA_03248 start codon. Notably, the Tn insertion site in the intergenic region, at 3415650, is only 188 bp away from the peak summit of one of the significant (although not top 5%) HigX binding sites, at 3415838, suggesting that this gene is a direct HigX target (Fig 6C). The two Tn mutants obtained from the pMT335-*higX* tolerance screen had Tn mutations in CCNA_00669, encoding a lipopolysaccharide biosynthesis protein, and approximately 100 bp upstream of the peptidoglycan amidase gene *amiC*. However, the two mutants obtained from the pMT335-*higX* tolerance screen were also ciprofloxacin-resistant (Fig. 6D), and the mutants obtained from the ciprofloxacin resistance screen could be transformed with pMT335-*higX*, demonstrating that even if the genetic screens were not saturated, the selection strategies have succeeded in isolating suppressor mutants with convergent phenotypes.

Taken together, these data support a model where overproduction of HigX in a Δ*lexA* background, either from endogenous overproduction in Δ*lexA* strains or *in trans* from pMT335-*higX*, leads to destabilization of the cell envelope. Since *higX* deletion protected the Δ*lexA* strain against Cipro and cephalexin sensitivity even though HigX did not bind its target genes properly in the Δ*lexA* mutant, the phenotype cannot be due to changes in HigX-dependent gene regulation but could be a property of simply having too much HigX protein in the cell membrane, or having the protein present but nonfunctional. We infer that our suppressor Tn mutants counteract the destabilizing effect of HigX on the cell envelope, either directly or indirectly.

### *higX* is widespread among alpha-proteobacteria but synteny with *higBA* is not conserved

To investigate whether this unusual combination of TA system with adjacent but independent transcription factor was conserved among other bacteria, we evaluated the genomic conservation of this locus using the FlaGs software (26) with the *Caulobacter crescentus* NA1000 HigX protein sequence as query and the limit of aligned genomes set to 99, of which 68 were successfully aligned. We observed that *higX* is not unique to *Caulobacter* species but found in many other alpha-proteobacteria as well. Within the *Caulobacter* genus, the flanking gene context of *higX* was well conserved, immediately upstream of the master cell cycle regulator *ctrA.* Moreover, it was co-conserved with a conserved uncharacterized gene, annotated as a putative hydrolase of the COG4188 family in all except five of the more distantly related genomes, including in the non-*Caulobacter* species which did not have their *higX* orthologue adjacent to *ctrA* (Fig. S2). However, in the 68 aligned *higX*-proximal regions, only four of the genomes contained probable *higBA* orthologues upstream of *higX*. Hence, *higX* is much more conserved in its genomic context in alpha-proteobacteria than *higBA* is (although we do not exclude that the other bacteria may encode *higBA* at a different site in the genome, since at least in the *Caulobacter* genus it is conserved between species (27)) and *higX* is almost universally found adjacent to the uncharacterized hydrolase gene.

While examining the alignments of Fig S2 we noted that the intergenic region between the putative hydrolase and the *higB* gene appeared consistently larger than the intergenic region between the putative hydrolase and the *higX* gene (in the majority of genomes which lacked *higBA*), suggesting that some of the intergenic sequence may be associated exclusively with the *higBA* genes. We therefore aligned the intergenic regions between the putative hydrolase gene and *hig(BA)X* from a subset of 15 *Caulobacter* genomes, including NA1000 and the four others which contained *higBA* upstream of *higX*, and observed a consistent breakpoint in the intergenic region where the intergenic sequence stopped in the genomes lacking *higBA*, but continued in the genomes which had *higBA* upstream of *higX*. (Fig. 7A). This breakpoint was at base 3281622 of the *Caulobacter crescentus* NA1000 genome, relative to the start codons of the putative hydrolase at base 3281744 and *higB* at base 3281524. From this, we inferred that of this intergenic region, the 122 bp adjacent to the putative hydrolase gene is universal to all the genomes, while the additional 98 bp of intergenic sequence found only in the genomes of the strains possessing *higBA* is likely to be a *higBA*-specific regulatory element. Notably, this additional sequence contained the high-affinity LexA binding site that is responsible for the LexA-mediated repression of the *higBA* promoter. Therefore, the LexA-mediated repression of *higX* under normal conditions and the overproduction of HigX when this repression is released is likely unique to NA1000 and the other *Caulobacter crescentus/vibrioides* strains which contain *higBA* and its LexA-dependent promoter immediately upstream of *higX*. In all other *higX*-encoding alphaproteobacteria, we would instead anticipate *higX* to be expressed in a LexA-independent manner.

**Figure 7.**
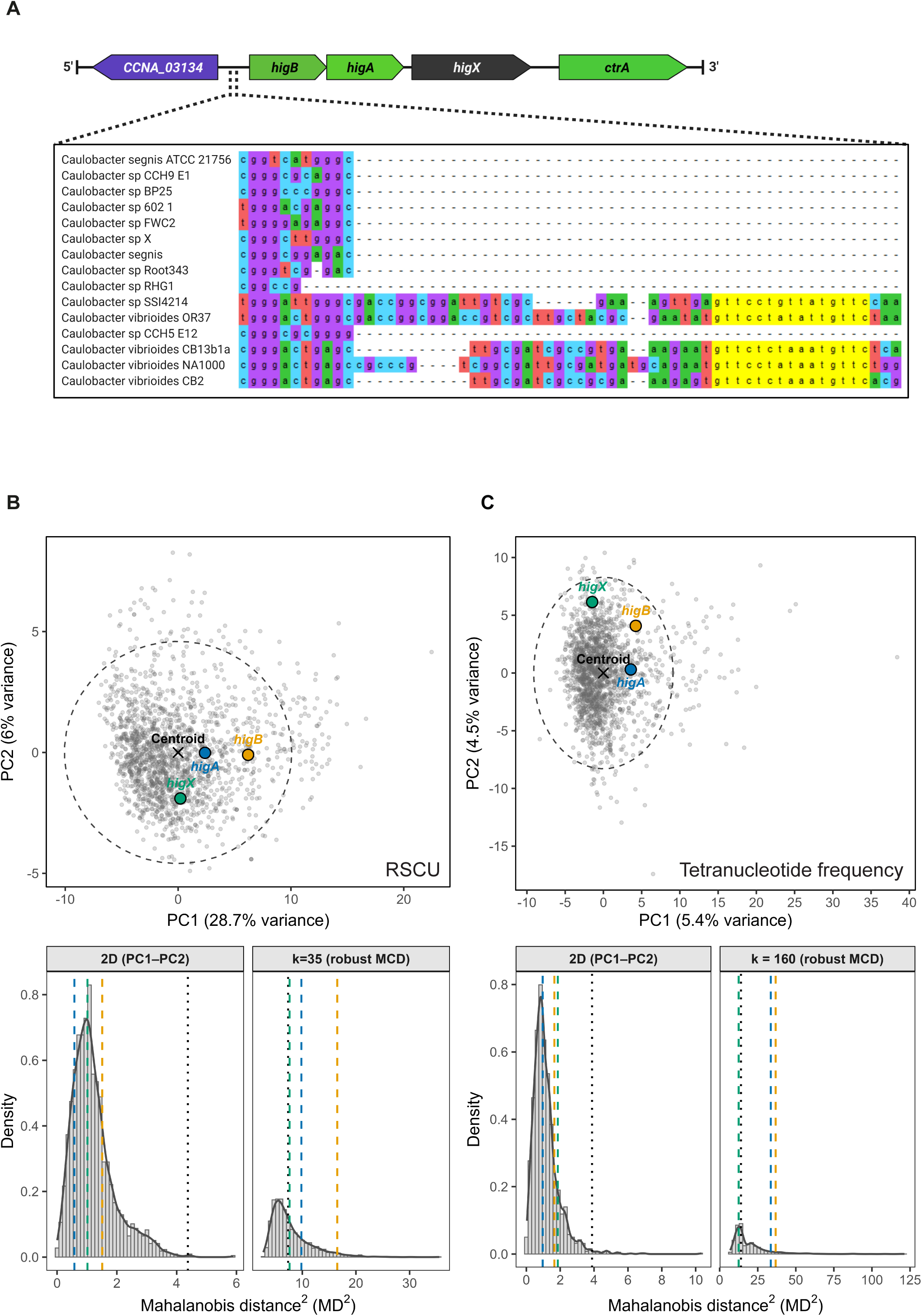
HigX is broadly conserved in its genomic context while higBA is restricted to the Caulobacter crescentus/vibrioides lineage. (A) Alignment of the intergenic region upstream of *hig(BA)X* containing the breakpoint between *+higBA* and *-higBA* strains. The DNA sequence shown corresponds to bases 3281574 to 3281632 of the NA1000 genome sequence. The LexA binding motif is shown in yellow. (B) Multivariate compositional analysis of relative synonymous codon usage (RSCU) across all coding sequences in *Caulobacter* NA1000 with individual CDSs represented as grey points and the 95 % confidence intervals around the genomic centroid indicated as a dashed ellipse. The three genes of interest are indicated as *higB* (orange), *higA* (blue) and *higX* (green). Below the PCA plot, density plots display the genome-wide distributions of Mahalanobis distance (MD²) under both standard two-dimensional (PC1–PC2) and robust minimum covariance determinant (MCD) models (k = 35). Empirical FDR < 0.05 cutoffs (Benjamini–Hochberg), are shown as black dotted lines and the MD^2^ values for *higB*, *higA* and *higX* are indicated in their respective colors. (C) Multivariate compositional analysis of tetranucleotide frequencies using the same approach as in (B). The tetranucleotide frequency composition of each gene was normalized and projected into principal component space, with the 95% confidence intervals around the genomic centroid indicated as a dashed ellipse. Once again, *higB*, *higA*, and *higX* are shown in their respective colors, and the density plots display the corresponding MD² distributions for both two-dimensional and robust MCD models (*k* = 160) with FDR-based significance thresholds.

In order to investigate whether *higBA* is a recent arrival in a minority of the *Caulobacter* genomes, or whether it is an ancestral factor that has been lost from the majority of them, we employed compositional analysis of relative synonymous codon usage (RSCU) and tetranucleotide frequency. Using multivariate compositional analysis of these two parameters (Supplementary Dataset S2), we detected a minor atypicality of *higX* in its codon usage (q = 0.0211), but no significant difference in tetranucleotide composition (q = 0.805) relative to other coding sequences of the genome. In contrast, *higB* and *higA* displayed significant compositional atypicality in their RSCU (*higB*: q = 3.83 x 10^-37^, *higA*: q = 5.25 x 10^-7^) and tetranucleotide composition (*higB*: q = 9.46 x 10^-187^, *higA:* q = 4.39 x 10^-144^), compared to all CDS within the genome. This degree of deviation from the rest of the genome is potentially indicative of horizontal gene transfer or, at the least, evidence for a recent arrival (Fig. 7B+C). The small deviation in codon usage for *higX* is likely biologically negligible, with an effect size of 1.08 (i.e., 8% above the 95% cutoff; Figure S3A). By comparison, *higB* and *higA* exhibited larger effect sizes (RSCU: 2.34 and 1.39; tetranucleotide: 2.66 and 2.43; Figure S3A). Thus, *higBA* appear to be compositionally highly atypical, whereas *higX* is compositionally typical relative to the rest of the genome. We conclude from this that the presence of *higBA* at this locus is most likely due to an insertion event into a recent ancestor of the NA1000 lab strain of *Caulobacter* and a few closely related species, while the presence of *higX* alone at this locus is the “ancestral” state.

### *higX* is ancestral but *higBA* is of likely foreign origin

To support the compositional bias analysis, we investigated if the compositional typicality of *higX* and atypicality of *higBA* was reflected in their encoded proteins’ phylogeny, as an indication of their most likely origin. The 16S rRNA phylogenetic tree showed that all genera cluster into separate clades, with the only exception being *Asticcacaulis excentricus* CB 48, whose placement is within the *Brevundimonas* clade (Fig. S3B). Regarding the presence/absence heatmap, as expected, HigX was found to be the most prevalent protein across the order, with 13 organisms containing it in different combinations and 9 containing it as the sole protein. Interestingly, the prevalence of all 3 proteins was not limited to members of the *Caulobacter* genus. In fact, one member of *Brevundimonas* contained all three proteins. However, it is important to note that this heatmap does not account for the synteny of the genes encoding the proteins. Thus, it is possible that even if an organism contains all three proteins, they are not within the same gene cluster or operon. *higBA* could likely be positioned away from *higX* somewhere else on the genome.

The protein phylogenetic trees to be compared against the 16S rRNA phylogenetic tree were generated by utilizing BLASTx with *higB, higA* or *higX* as queries. Hits (n =100) were filtered and used for creating a presence/absence heatmap of each gene per organism and in different combination classes (Fig S4). Occurrence of all three proteins was limited to members of the *Caulobacter* genus (n = 5), closely matching results of the heatmap from the 16S rRNA phylogenetic tree (Figure S3B). HigB and HigA co-occurring in the absence of HigX were only found in one species out of the 100 used and the co-occurrence of HigA and HigX was limited to the *Caulobacter* genus (n = 7). Interestingly, in the remaining combination classes, a stark contrast was observed. The class for HigB as the sole protein (n = 41) contained no organisms within the Caulobacterales order (Fig S4, S5). In fact, twelve of these were members of the *Enterobacter* genus, which is part of Gammaproteobacteria, showing a significant discrepancy with the 16S rRNA phylogenetic tree. A similar pattern was observed for the HigA-only class (n = 59), whose members consisted of organisms outside Caulobacterales (Figure S4, S6). By comparison, the HigX-only class (n = 56) displayed a phylogeny similar to the 16S rRNA phylogenetic tree, with only two organisms coming from outside the Caulobacterales order (Figure S4, S7). Overall, the results from the phylogenetic analysis strongly indicate that *higX* is both ancestral to and phylogenetically consistent with the Caulobacterales lineage, whereas the phylogenies of *higB* and *higA* are indicative of a foreign genetic origin, possibly enterobacterial in the case of *higB*.

## Discussion

In this study, we identify HigX as the factor responsible for the unexpected and HigB-independent ciprofloxacin sensitivity of the Δ*lexA* Δ*higBA* mutant. Although HigX appeared at first to be a third component of the HigBA TA system, due to its close genetic proximity, annotation as transcription factor and ability to weakly regulate the *higBAX* promoter, we found that it does not exert its effect through the TA system. Instead, it acts independently at the level of the cell envelope where it is physically located and for which it regulates some of the genes. Δ*lexA* mutant strains produce higher levels of HigX due to the placement of the *higX* gene directly downstream of the strong LexA-dependent promoter of *higBA* and the lack of LexA to repress it, but this is not reflected in increased binding of HigX to its target promoters. HigX is incorporated into the cell membrane through its N-terminal domain, and high levels of HigX sensitize Δ*lexA* cells, but not WT cells, to ciprofloxacin and cephalexin. Forward genetic screening showed that toxicity of excessive HigX production can be suppressed by cell envelope-related mutations, including of at least one HigX target gene.

Our compositional bias and phylogenetic analysis of *higBAX* revealed that higX appears to be ancestral and phylogenetically conserved with a strong match to the 16S rRNA phylogeny of *Caulobacter*. In contrast, *higB* and *higA* were found to be atypical in both the compositional parameters (RSCU and tetranucleotide) and their phylogeny, suggesting a foreign nature. Unexpectedly, the phylogeny of HigB was very different compared to HigA. The genus *Enterobacter* was prevalent in the HigB tree (n=13), thus indicating this genus as the most likely origin of *higB.* However, the potential origin of *higA* is less clear, as its prevalence is less clustered to a single genus like *higB* and it did not display the same bias towards enterobacterial orthologues. Instead, its occurrence is highest among *Stenotrophomonas* (n=10), *Bradyrhizobium* (n=7), and *Xanthomonas* (n=5). Therefore, the potential source of *higA* acquisition can be reduced to these three genera, with *Stenotrophomonas* being the most likely candidate. Taken together, these results reveal that the toxin and antitoxin are of different evolutionary lineages and were most likely acquired separately, rather than together, as previously documented (reviewed in (9, 28)).

Our initial observation that led us to investigate *higX* was the ciprofloxacin sensitivity phenotype of the Δ*lexA* mutant. This had initially seemed counter-intuitive, given that this strain has the SOS response constitutively activated in defence against DNA damaging drugs. However, part of the SOS response in *Caulobacter* is to inhibit cell division through the divisome inhibitor protein SidA (29) leading to the production of highly filamentous cells. *Caulobacter crescentus* is very sensitive to cell envelope stress relative to other Gram-negative bacteria (30) and it could be that extreme cell elongation weakens the integrity of the cell envelope, providing easier access to the cytoplasm for antibiotics and accounting for the ciprofloxacin sensitivity. Excessive incorporation of cell envelope proteins into the membrane and periplasm has been shown to cause cell envelope stress in elongated cells where the filamentation was caused by different factors than the SOS response (20), showing that LexA is not required for this effect. We propose that cell elongation-dependent weakening of the envelope could be the reason why HigX overexpression is toxic to Δ*lexA* cells, while WT cells are unaffected.

It was striking that global DNA binding of HigX was decreased in the Δ*lexA* cells even though they produced more HigX protein than WT cells did. We find it unlikely that the reduced DNA binding of HigX in the Δ*lexA* mutant is due to a global dependence on LexA for HigX binding, because in comparing our regulon of HigX to the previously established ChIP-Seq regulon of LexA (15) we found only 13 shared peaks, none of which were in our top 5% 30 HigX peaks. This suggests that access of HigX to its target promoters is the major limiting factor in the Δ*lexA* strain. Restriction of the protein to the inner face of the membrane would not limit nucleoid access in WT cells, because the chromosomal DNA in *Caulobacter* does not condense but fills the whole cell during the normal replication cycle (31). However, in filamentous cells, the DNA content is variable and depends on whether chromosome replication has arrested or continued (32). If replication continues, the DNA can continue to fill the cell volume, at least up to a point (33), but if it is inhibited then the filamentous cell will have one or two nucleoids surrounded by DNA-free regions (34, 35). The nucleoid arrangement in *Caulobacter* cells lacking LexA has not been imaged directly. However, the DNA content of Δ*lexA* cells as measured by FACS is characteristic of replication arrest with one or two chromosomes without over-replication (15), similar to cells with the SOS response induced by mitomycin C (32). This would be insufficient, in most cases, to fill up the entire cytoplasm of the highly filamentous Δ*lexA* cells. Therefore, even if more HigX is produced in Δ*lexA* cells, some of the available membrane surface into which it can be inserted could be away from the nucleoid, sequestering the protein away from the DNA and explaining the globally reduced binding that we observe. Removal of HigX from its membrane localization does not improve DNA access, since in our experiments with the truncated HigX lacking the four TM helices, we saw neither gene regulation activity nor GFP fluorescence suggesting that the TM helix bundle is required for the protein’s stability. Together, these data support a model where correct HigX regulatory activity can only happen with membrane-localised WT protein in normal shaped cells, while activity is lost in elongated cells or if the HigX TM domain is missing (Fig 8). It will be interesting to investigate whether HigX-dependent gene regulation is affected in cells that are filamentous for other reasons (independent of LexA) in order to test this hypothesis, as this would be more representative of HigX-dependent gene regulation in species that do not have the *higBA* TA system genes and their LexA box immediately upstream.

**Figure 8.**
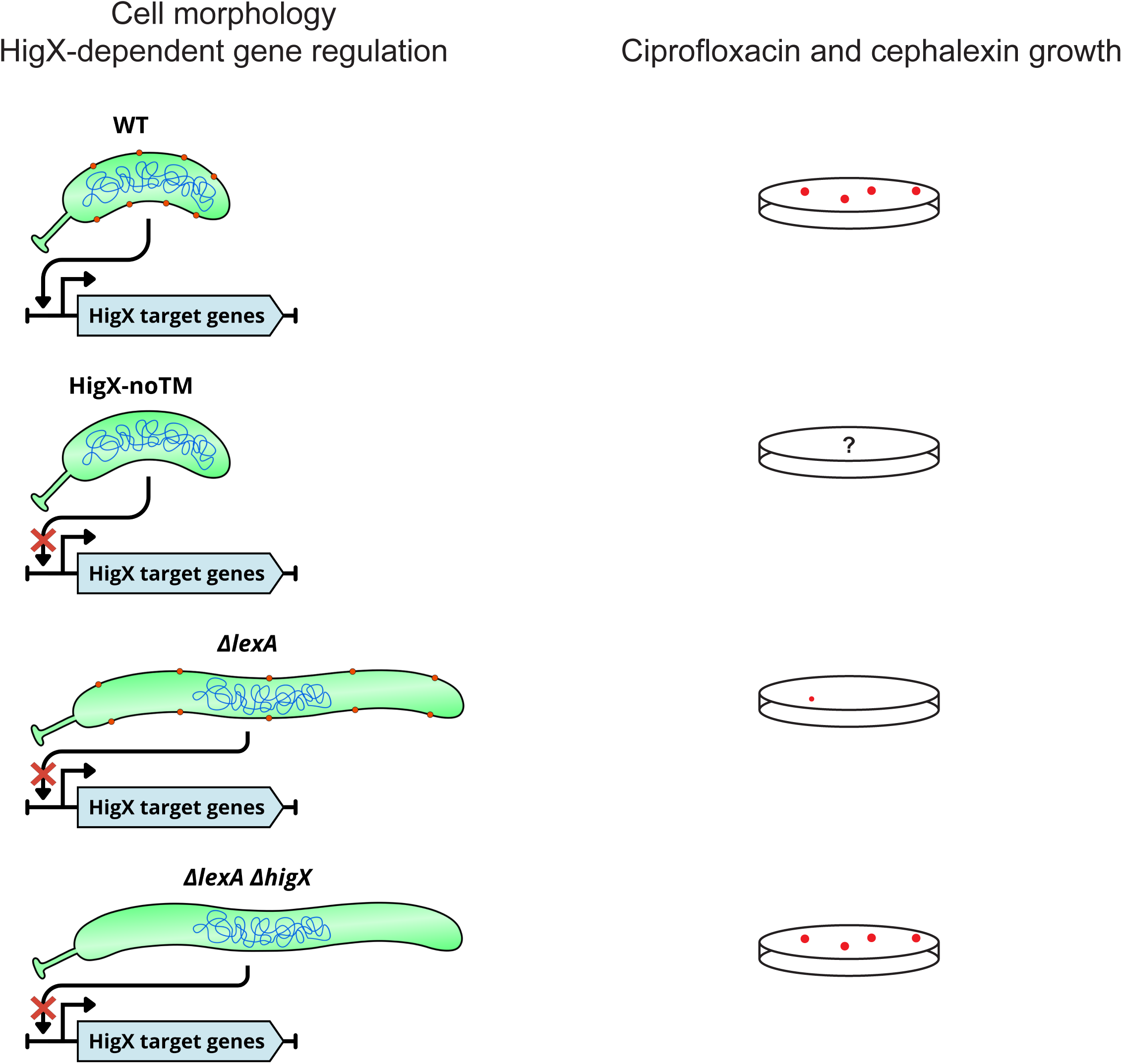
Model for the cell-shape dependent regulatory activity of HigX in Caulobacter. HigX promoter binding is active in WT cells with native HigX protein, but not in filamentous Δ*lexA* cells or WT shaped cells expressing HigX without its N-terminal transmembrane (TM) domain which is likely unstable or inactive. The nucleoid is represented in blue and HigX in orange. Resistance to ciprofloxacin and cephalexin is correlated with loss of HigX itself rather than loss of HigX DNA-binding activity (? indicates that this has not been tested for HigX-noTM).

Analysis of the regulon of HigX suggests that its cell membrane localisation may be linked to its regulation of processes that happen there. Indeed, the highest peak that we observed was associated with a periplasmic multidrug efflux lipoprotein precursor gene (CCNA_03218) (Fig 5B), similar to the AcrA component of AcrAB-NodT family multidrug efflux pumps. The COG analysis of the 5% top peaks had a noticeable representation of genes associated with transport of carbohydrates, transport of inorganic ions, energy production/conversion and cell envelope or membrane biogenesis (Fig 5C). We have not yet observed any phenotype linked to loss of HigX in WT cells grown under lab conditions. However, its role may be more important for envelope maintenance under stress conditions. The high level of evolutionary conservation of HigX across the Caulobacterales order suggests that its presence is generally beneficial, even if currently mysterious.

HigX’s transmembrane conformation is unusual for a LytTR family protein. The majority of these transcription factors are cytoplasmic and regulate extracellular processes such as competence, alginate production, expression of virulence factors and bacteriocins (23). However, little is known about how they are regulated themselves. In the original bioinformatic study which characterized the LytTR transcription factors, it was noted that all of the instances of this domain discovered in alpha-proteobacteria were probable membrane proteins (23), but without apparent periplasmic domains that could allow them to function as extracellular sensors in the same way as the membrane-bound transcription factors of *Vibrio cholerae* (36). Instead, their domain structure is more similar to the enterococcal bacitracin-sensing response regulator BcrR, which is permanently anchored to the inner membrane by a 4 helix transmembrane domain that senses bacitracin directly and stimulates transcription of target genes by the DNA-binding domain (37). More recent studies have identified a LytTR sub-family where the transcription factor is negatively regulated by a separate membrane protein, designated “LytTR regulatory systems” (LRS). First identified in *Streptococcus mutans* (38, 39), LRS are particularly abundant in the Firmicutes but also found in some actinobacteria and a few gamma-, beta-and epsilon-proteobacteria (40). In LRS the LytTR protein is consistently encoded upstream of the membrane protein, which usually consists of three or four transmembrane alpha-helices and no apparent periplasmic domain, and the membrane protein negatively regulates the transcription factor through membrane sequestration preventing it from activating its target genes (41). The similarity of the topology of LRS membrane proteins to the N-terminus of HigX (and the other Caulobacter LytTR proteins) is suggestive of the alpha-proteobacteria having developed a new family of “fused LRS” where the transmembrane domain and the DNA binding domain are in the same protein. However, the regulatory influence of the transmembrane domain on the DNA binding domain would have to be positive rather than negative, since we observe that full length HigX can carry out promoter regulation while the HigX DNA binding domain without the transmembrane domain cannot (Fig. 3D), similarly to BcrR where the DNA binding domain has no affinity for DNA without the transmembrane domain (37). A recent computational study has revealed that transmembrane transcription factors are widely prevalent throughout bacterial and archaeal taxonomic groups and may use the biophysical state of the membrane as a signaling input to regulate gene expression (42), which could explain the dependence that we see on the presence of the transmembrane domain for HigX.

In summary, we have uncovered a role for the first of this class of transmembrane transcription factors from the alpha-proteobacteria to be characterized, and shown that cell filamentation through loss of *lexA* negatively affects the ability of HigX to bind its target genes. In addition to identifying a new type of cell envelope regulatory factor in *Caulobacter*, our findings also indicate that TA system incorporation into a bacterial genome can cause alternative downstream phenotypes that do not depend on the function of the TA system itself, but on the regulatory element(s) that it brings along and the genomic context into which it arrives.

## Methods

### General growth conditions

*Caulobacter crescentus* strains were routinely grown in peptone-yeast extract (PYE) medium at 30°C and *E. coli* strains in LB at 37°C. Antibiotics were used at the following concentrations; tetracycline at 1 μg/ml for *Caulobacter* and 10 μg/ml for *E. coli*, gentamicin at 1 μg/ml for *Caulobacter* and 10 μg/ml for *E. coli*, nalidixic acid at 20 μg/ml for *Caulobacter* and kanamycin at 20 μg/ml (solid media) or 5 μg/ml (liquid media) for *Caulobacter* and 20 μg/ml for *E. coli*. Ciprofloxacin was prepared as 20 mg/ml stock solution in 0.1 M HCl and in all experiments involving ciprofloxacin, an equivalent volume of 0.1 M HCl was added to the control cultures or plates. Vanillate stock solution was prepared at 50 mM stock solution (adjusted to pH 8.0 with NaOH) and used at 50 or 100 μM as indicated in the figure legends.

### Strain and plasmid construction

DNA fragments for cloning were PCR-amplified with Phusion DNA polymerase (Thermo Fisher Scientific) from stationary phase cultures of wild type (WT) *Caulobacter crescentus*, using PCR primers listed in Supplementary Table S2. Products were purified by agarose gel electrophoresis. Cloning of the correct region was confirmed by sequencing and plasmid stocks were maintained in *E. coli* EC100 or TOP10. Plasmids used in this work are listed in Supplementary Table S3. Replicating plasmids (pMT335 (43), pJC327 (44) and plac290 derivatives) were transferred into *Caulobacter* strains by electroporation while suicide plasmids for generation of deletion mutants (pNPTS138 derivatives) and transposon mutants (pHPV414 (45)) were transferred by conjugation from *E. coli* S17-1 *λpir* (46). After integration of suicide vectors by recombination through one of the homologous flanking regions, secondary recombination events were induced by counterselection on PYE agar containing 3% sucrose and mutants carrying resulting in-frame deletions were screened for by PCR. Double mutants were made by introducing the Δ*higX* or Δ*higBAX* alleles into the Δ*lexA* mutant strain. Generalised transductions with bacteriophage ΦCr30 were performed as previously described (47). *C. crescentus* strain numbers and genotypes are listed in Supplementary Table S4. pET28a derivatives containing WT or truncated HigX fused to an N-terminal His-tag were expressed in *E. coli* Rosetta(DE3) (Novagen).

### Efficiency of plating assays

Resistance to cephalexin and ciprofloxacin was assessed by dilution spot plating. Cultures of strains to be tested were grown overnight to stationary phase, then inoculated into new medium to grow to mid-exponential phase. Culture density was measured, normalised to the OD600 of the least dense culture (OD600=0.5 or less), serially diluted in PYE to 10^-6^ and 5 µl spotted onto plates containing PYE medium with sub-inhibitory concentrations of cephalexin (5 μg/ml) or ciprofloxacin (1 μg/ml). For *higX* overexpression experiments, the plates additionally contained gentamicin in order to maintain selection on the plasmid, and all plates contained vanillate (50 μM) to induce overexpression of *higX*. Plates were imaged after 3 days growth at 30°C. Images are representative of at least three independent biological replicates.

### Beta-galactosidase assays

β-galactosidase assays were performed on strains carrying low copy plasmid-borne transcriptional or translational fusions of the promoters of interest to l*acZ*. Cultures were grown to early exponential phase (OD_600_ = 0.1 – 0.4) with exposure to ciprofloxacin or vanillate as described in the main text or figure legends, followed by β-galactosidase assays using the method of Miller (35) on three independent biological replicates.

### RNA extraction and quantitative RT-PCR

RNA was extracted from 4 ml mid-exponential phase cultures which were treated with 2500 U Ready-Lyse (Bionordika) and homogenized using QiaShredder columns, prior to RNA extraction with the RNEasy Mini Kit (Qiagen) according to the manufacturers’ instructions, including on-column DNase digestion. RNA quality was assessed by agarose gel electrophoresis and concentration was measured in a Nanodrop spectrophotometer. cDNA was prepared using the SuperScript IV Reverse Transcriptase Kit (Thermo Fisher) according to the manufacturers’ instructions, on 400 ng RNA template using random hexamer primers. Quantitative RT-PCR was performed in technical and biological triplicates on a 96-well LightCycler real-time PCR system (Roche) using SYBR Green (Roche). The *higB* transcript was amplified with primers higB_qrt_fwd and higB_qrt_rev, the *higX* transcript was amplified with primers CC_3036_qrt_fwd and CC_3036_qrt_rev, and the reference gene *rpoD* was amplified with primers rpoDfow and rpoDrev. Quantification was by the standard curve method and *higB* and *higX* transcript levels were normalized to *rpoD*.

### Western blot and anti-HigX antibody

Custom polyclonal antibody to HigX for western blotting and ChIP-Seq was developed by Nordic Biosite. Rabbits were immunized with the synthetic peptide (C-)LRMEDHYVRIRTEHGSRLE corresponding to amino acids 178 to 196 of the HigX protein with cysteine conjugated at the N-terminus. Antiserum was extracted from the rabbits and affinity purified on a Sepharose column containing the immobilized synthetic peptide to obtain purified polyclonal antibody. Cell extracts for western blotting were prepared by growing *Caulobacter* cultures in PYE (containing 50 µM vanillate for strains containing pMT335 derivatives), or *E. coli* Rosetta(DE3) cultures in LB with 0.1 mM IPTG, to mid-exponential phase, normalizing to OD600 = 0.5, centrifuging and resuspending in 100 µl SDS-PAGE loading buffer. Extracts were run on a 16% SDS-PAGE gel alongside PageRuler prestained markers (Thermo Fisher). Proteins were transferred to PVDF membranes and blocked by incubation in TBS containing 5% dried skim milk. HigX protein was detected by incubation with primary antibody diluted 1/5000 in TBS with 2% skim milk at 4°C overnight, secondary antibody diluted 1/4000 in TBS with 2% skim milk at 4°C for one hour, and HRP substrate at room temperature for one minute. Between incubations, membranes were washed 2 x 5 minutes then 1 x 15 minutes in TBS.

### Microscopy

Cultures for imaging were grown to mid-exponential phase with 50 μM vanillate to induce HigX-GFP expression and immobilized on 1% agarose (in water) microscope slides. Phase-contrast and GFP fluorescence images were obtained with an inverted Olympus IX83 microscope equipped with a 100x/1.46 oil objective and a Photometrics Prime scientific complementary metal-oxide-semiconductor (sCOMS) camera. GFP images were taken before phase-contrast images to minimize bleaching. Images are representative of three independent biological replicates.

### Quantitative mass spectrometry

Cultures for mass spectrometry were grown to mid-exponential phase, harvested by centrifugation and cell pellets were immediately frozen in liquid nitrogen. Cells were sonicated in ice-cold 100 mM Na_2_CO_3_ containing cOmplete protease inhibitor and PhosSTOP phosphatase inhibitor (Sigma) for 2×20 sec at 60% amplitude on ice using a probe sonicator. After sonication the samples were incubated at 4 °C for 1h with rotation. After incubation, the samples were ultracentrifuged at 100.000g for 90 min using a Hitachi Micro ultracentrifuge (CS 150 NX). After ultracentrifugation, the supernatants (soluble protein fraction) were transferred to another tube and the membrane pellets were gently washed with 100µL 50 mM TEAB, pH 8.5. The membrane pellets were resolubilized in 5% sodium deoxycholate (SDC) in 50 mM HEPES buffer, pH 8.5, by probe sonication (1×20 sec at 20% amplitude). After sonication, the samples were centrifuged at 20.000g for 10 min, the supernatants were transferred to new tubes and the protein concentrations were measured by Nanodrop (Implen Nanophotometer N60). The membrane protein samples were reduced and alkylated using 5 mM DTT for 30 min and 10 mM iodoacetamide for 30 min, respectively. After alkylation, trypsin was added (5%) and the samples were incubated at 37 °C over-night. After incubation, additional trypsin (1%) was added and incubated at 37 °C for 1 hour.

A total of 25 µg of peptides from all conditions were labeled with Tandem Mass Tag (TMT) pro 18 plex (Thermo Fisher Scientific) according to the manufacturer’s protocol. The TMT labeling was performed as following: NA1000 (TMT 126, 127N and 127C); *ΔhigBA* (TMT 128N, 128C and 129N); *ΔhigX* (TMT 129C, 130N and 130C); *ΔlexA* (TMT 131N, 131C and 132N); *ΔlexA ΔhigBA* (TMT 132C, 133N and 133C). After labeling, all samples were combined into one sample and the SDC was precipitated by adding formic acid to 2% followed by centrifugation at 20.000g for 20 min. The supernatant was transferred to a new Eppendorf tube and dried by vacuum centrifugation. The dried TMT labeled peptides were desalted using Poros Oligo R3 Reversed Phase (RP) material and subsequently fractionated using high pH RP fractionation into 12 concatenated fractions as described previously (PMID: 35240313). Each fraction was dried and resolubilized in 10 µL 0.1% formic acid prior to liquid chromatography tandem mass spectrometry (LC-MS/MS).

An aliquot of each peptide fraction was loaded onto a 20 cm analytical column (100 μm inner diameter) packed with ReproSil – Pur C18 AQ 1.9 μm RP material using an EASY nanoLC system coupled to an Orbitrap Eclipse Tribrid (Thermo Fisher Scientific). The peptides were eluted with an organic solvent gradient from 100% phase A (0.1% FA) to 22% phase B (95% ACN, 0.1% FA) for 100 min, then from 22% B to 40% B for 20 min before the column was washed with 95% B. The flow rate was set to 300 nL/min during elution. The automatic gain control target value was 1.5×10^6^ ions in MS and a maximum fill time of 50 ms was used. Each MS scan was acquired at high-resolution (120,000 full width half maximum (FWHM) at m/z 200) in the Orbitrap with a mass range of 350-1500 Da. The instrument was set to select as many precursor ions as possible in 3 sec between the MS scans. Peptide ions were selected from the MS for higher energy collision-induced dissociation (HCD) fragmentation (collision energy: 34%). Fragment ions were detected in the orbitrap at high resolution (50,000 FWHM) for a target value of 1,5×10^5^ ions and a maximum injection time of 150 ms using an isolation window of 0.7 Da and a dynamic exclusion of 20 sec. All raw data were viewed in Xcalibur v3.0 (Thermo Fisher Scientific).

All LC-MS/MS raw data files were searched in Proteome Discoverer (PD) version 2.5.0.400 (Thermo Fisher Scientific) using the SEQUEST HT search algorithm against the *Caulobacter crescentus* (strain NA1000 / CB15N) OX=565050 protein FASTA database. The searches had the following criteria: enzyme, trypsin; maximum missed cleavages, 2; fixed modifications, TMTpro (N-terminal), TMTpro (K) and Carbamidomethyl (C).

The TMTpro reporter ion signals were quantified using S/N and they were normalized to the total peptide S/N in the PD program. The in-built ANOVA test in PD was used to generate p-values for all the proteins identified in the database searches with 1% FDR.

### ChIP-Seq sample preparation and sequencing

Chromatin immunoprecipitation (ChIP) was performed essentially as described in Fumeaux et al (48) with the following modifications. Cultures were grown to mid-exponential phase in 80 ml PYE before formaldehyde crosslinking. Sonication was carried out in a Covaris ultrasonicator with the following settings: temperature 6°C, duration 3600 seconds, peak power 75.0, duty % factor 10.0, cycles per burst 200, average power 7.5, delay 60 seconds. All of the cleared sonicated lysate was used for ChIP without taking an input sample out first. HigX-bound DNA was recovered by immunoprecipitating with 20 µg purified anti-HigX antibody per sample, divided between two Eppendorf tubes. After immune complex recovery, reversal of crosslinks and protease treatment, DNA was recovered in 50 µl MilliQ water per Eppendorf and pooled into 100 µl, of which 40 µl was used for sequencing. DNA was prepared for Illumina sequencing using the NEBNext UltraExpress DNA library preparation kit (New England Biolabs) according to the manufacturer’s instructions and 50 bp paired-end sequencing was carried out on a NovaSeq 6000 (Illumina). Details of the sequence processing, alignment, quantification and data analysis are in the Supplementary methods.

### Transposon mutagenesis, forward genetic screening and insertion mapping

A pooled library of transposon mutants was constructed in the Δ*lexA* Δ*higBA* background by performing five independent conjugations of the transposon carrier plasmid pHPV414 from *E. coli* S17-1 *λpir* into the Δ*lexA* Δ*higBA* recipient strain. Transconjugant colonies were selected on PYE plates with nalidixic acid and kanamycin after three days incubation at 30°C. Colony counting indicated that the library consisted of > 25’000 independent clones. Colonies were pooled in PYE and stored in aliquots at-80°C in 10% DMSO.

The ciprofloxacin resistance screen was performed by growing the pooled library to stationary phase and diluting it into medium containing 1 μg/ml ciprofloxacin and 20 ug/ml kanamycin, followed by plating on PYE plates containing 1ug/ml ciprofloxacin and 20 ug/ml kanamycin, and incubation at 30°C for five days. 40 colonies were re-streaked on PYE plates containing 1 μg/ml ciprofloxacin and 20 ug/ml kanamycin followed by incubation for 4 days to confirm growth.

The HigX overexpression tolerance screen was performed by growing the pooled library to stationary phase, preparing electrocompetent cells and electroporating with pMT335-*higX*. Plasmid-tolerating colonies were selected by plating electroporated cells on PYE plates containing gentamicin and 50 μM vanillate. Colonies were re-streaked to new gentamicin/vanillate plates to confirm stability of the phenotype.

Transposon insertion sites were mapped by plasmid rescue. Genomic DNA was extracted from the ciprofloxacin-resistant or HigX-tolerant transposon mutants, partially digested with *Sau*IIIA and ligated. Ligations were used to transform *E. coli* EC100D, in which the transposon can be maintained as a replicating plasmid, and selected for kanamycin resistance. Plasmids were extracted and sequenced with the transposon-specific himup2 primer across the junction between the transposon insertion site and the flanking genomic DNA. To confirm specificity of the transposon insertions for the observed phenotypes, the transposon insertions were back-crossed into the Δ*lexA* Δ*higBA* strain by bacteriophage ΦCr30-mediated generalized transduction. The resulting strains were tested for both ciprofloxacin resistance and tolerance of electroporation with pMT335-*higX*.

### Sequence alignment and analysis

The genomic context conservation of HigX orthologues was investigated with FlaGs (26). Transmembrane segment prediction was performed with the DAS TM prediction program (49) and the structure was modelled in Alphafold using default parameters (50). Multiple sequence alignment of the region spanning *CCNA_03134* to *ctrA* was performed with MAFFT (51) (default parameters) and visualized with Molecular Evolutionary Genetics Analysis version 11 (52) (MEGA version 11.0.13). Details of compositional bias analysis, comparative genomics and phylogenetic analysis of proteins and 16S RNA sequences are in the Supplementary Methods.

### Statistical analysis

All numerical data are reported as mean of all biological replicates performed and error bars indicate the standard deviation. Statistical significance was analysed by non-paired equal variance 2-tailed Student’s T test for simple pairwise comparisons between strains or treatment conditions and by a Dunnett’s test in R (version 4.2.2) for multiple comparisons. Dunnett’s test in R was carried out by setting the wild-type strain (NA1000) as a reference and comparing all other strains against it. * signifies p < 0.05, ** signifies p < 0.01 and *** signifies p < 0.001 throughout.

## Supporting information

ChIP-Seq peak data file

Compositional analysis of higB, higA and higX

## Acknowledgements

This work was supported by a grant from Independent Research Fund Denmark and start-up funding from the University of Southern Denmark, both to CLK. We are grateful to Asmus Cosmos Skovgaard for technical assistance. We thank Bert Ely, Gemma Atkinson, Nathan Fraikin and Ditlev Brodersen for helpful discussions and Vishnu Narayanan Madhavan for critical reading of the manuscript. Sequencing of DNA for ChIP-Seq was carried out at the Center for Functional Genomics and Tissue Plasticity, Functional Genomics & Metabolism Research Unit, University of Southern Denmark. The authors thank Ronni Nielsen for sequencing assistance.

## Author contributions

CLK conceptualized and designed the study. KAB, NVH, KG, SN, LHH, LAJ, MRL and CLK performed experiments. MRL analysed the quantitative proteomics data. NVH and CLK analysed ChIP-Seq data and performed statistical analysis. KAB, KG, NVH, MRL and CLK wrote the paper.

## Data availability

The Fastq files containing the raw data from the ChIP-Seq experiments in WT and Δ*lexA* for HigX genome binding have been deposited in GEO under the accession number GSE310922. All other data is available within the paper or in its Supporting Information.

## Supplementary Figure legends

**Figure S1. Structure prediction of HigX as a transmembrane LytTR-family transcription factor.**

(A) Prediction of the transmembrane helix locations by DAS. (B) Model structure prediction of HigX by Alphafold, showing the N-terminal domain as a tightly packed four-helix bundle and the C-terminal domain with the typical LytTR DNA binding domain.

**Figure S2. Extended FLaGs analysis of genomic context around HigX shows that it is widely conserved with a divergent putative hydrolase enzyme across other alpha-proteobacterial genera.**

Output of FLaGs analysis using the results of a BLASTP search of HigX against known alpha-proteobacterial genomes (the BLASTP query sequence of *Caulobacter crescentus* NA1000 HigX is not included here, only the BLASTP output), with the limit of aligned genomic regions increased to 99, of which 68 were aligned. The number codes for selected proteins with annotated functions are as follows: 1, uncharacterized putative hydrolase; 2, FliI; 3, CtrA; 4, FliJ; 5, LdpD; 22, HigA; 25, HigB.

**Figure S3. Effect size of the compositional bias and phylogenetic conservation of the higBAX locus.**

(A) Effect size analysis of *higB*, *higA*, and *higX* based on codon usage (RSCU) and tetranucleotide frequency composition. Bars represent the square root of the Mahalanobis distance^2^ (√MD^2^) relative to the 95% robust cutoff (MCD estimator), expressed as a fold over the threshold. The 95% cutoff (effect size = 1) is indicated as a dashed horizontal line, meaning that values above this indicate deviation from the genome-wide compositional background. (B) Phylogenetic tree of the order Caulobacterales showing the distribution of the proteins HigB, HigA, and HigX. The maximum-likelihood tree was constructed from 16S rRNA sequences of complete and reference genomes (n = 24) retrieved from NCBI, serving as a phylogenetic standard for comparison. The accompanying heatmap indicates the presence or absence of the three proteins across the same genomes. Note that the heatmap does not account for gene synteny.

**Figure S4. Distribution of HigB, HigA, and HigX homologs across bacteria**

Presence–absence heatmap showing the occurrence of HigB, HigA, and HigX homologs across bacterial genomes identified by BLASTX searches against the NCBI RefSeq protein database. Each column indicates one of the three proteins, and each row corresponds to an organism with both its scientific name and NCBI TaxID. Presence/absence (blue/grey) is indicated by the left sidebar, and the right sidebar indicates protein combinations. Note that the heatmap does not account for gene synteny.

**Figure S5. Phylogeny of HigB homologs across bacteria**

Phylogenetic tree of HigB homologs based on protein sequences retrieved from the top BLASTX hits. Sequences were aligned with MAFFT (L-INS-i), followed by inference of the tree with IQ-TREE using the maximum-likelihood method. The tree was midpoint-rooted for visualization, with tip labels displaying organism names, which have been colored by genus. Bootstrap support values (≥ 50) are displayed at internal nodes, and the scale bar indicates the number of amino-acid substitutions per site. The phylogram in the upper left corner shows a condensed version of the 16S phylogenetic tree (Figure S4B), which acts as a phylogenetic standard for comparison.

**Figure S6. Phylogeny of HigA homologs across bacteria**

Phylogenetic tree of HigA homologs based on protein sequences retrieved from the top BLASTX hits. Sequences were aligned with MAFFT (L-INS-i), followed by inference of the tree with IQ-TREE using the maximum-likelihood method. The tree was midpoint-rooted for visualization, with tip labels displaying organism names, which have been colored by genus. Bootstrap support values (≥ 50) are displayed at internal nodes, and the scale bar indicates the number of amino-acid substitutions per site. The phylogram in the upper left corner shows a condensed version of the 16S phylogenetic tree (Figure S4B), which acts as a phylogenetic standard for comparison.

**Figure S7. Phylogeny of HigX homologs across bacteria**

Phylogenetic tree of HigX homologs based on protein sequences retrieved from the top BLASTX hits. Sequences were aligned with MAFFT (L-INS-i), followed by inference of the tree with IQ-TREE using the maximum-likelihood method. The tree was midpoint-rooted for visualization, with tip labels displaying organism names, which have been colored by genus. Bootstrap support values (≥ 50) are displayed at internal nodes, and the scale bar indicates the number of amino-acid substitutions per site. The phylogram in the upper left corner shows a condensed version of the 16S phylogenetic tree (full version in Figure S4B), which acts as a phylogenetic standard for comparison.

