## Supplemental Methods and Supplemental Tables S1 to S4 for "An ancestral transmembrane transcription factor couples cell envelope regulation and the SOS response in *Caulobacter crescentus*"

Contains Supplementary Methods text and four supplementary tables.

**Supplementary Methods**

***ChIP-seq data analysis***

Fastq files of the raw Illumina sequencing data were uploaded to the Galaxy server (1) for pre-alignment processing, alignment to the genome, quality control and peak calling. The Cutadapt tool was used in paired-end mode to trim the adapter sequences (Illumina TruSeq) AGATCGGAAGAGCACACGTCTGAACTCCAGTCA from Read1 and AGATCGGAAGAGCGTCGTGTAGGGAAAGAGTGT from Read2 sequencing reads. The trimmed reads were aligned to the reference genome (accession number NC_011916.1) using the bowtie2 tool in paired-end mode with default alignment settings. The BAM files obtained from bowtie2 were quality filtered using the “filter SAM or BAM, output SAM or BAM” tool so that only alignments with a MAPQ quality score of ≥ 30 were kept.

Filtered BAM files were then subjected to peak calling by utilizing the “MACS2 callpeak” tool. All settings were kept as default except for the following parameters. “Format of Input Files” was set to “Paired-end BAM”, “Effective genome size” was set to “User defined”, and the exact size of the *C. vibrioides* NA1000 genome was used (4.042.929 bp). For “Build model”, the setting “Do not build the shifting model (--nomodel)” was selected. Lastly, “Peaks as tabular file”, “Peak summits”, and “Summary page (html)” were chosen as additional outputs.

The quality of the ChIP-seq data was then assessed based on three quality metrics. Fraction of reads in peaks (FRiP), normalized strand cross-correlation (NSC), and relative strand cross-correlation (RSC). The FRiP score calculation was performed in R version 4.5.0 (2025-04-11) using RStudio 2024.12.1+563 (“Kousa Dogwood”). Specifically, the paired-end reads from the filtered BAM files for peak calling were collapsed into fragments and filtered by retaining properly paired alignments and removing unmapped, secondary, supplementary, and duplicate reads. Moreover, each read in the pair was filtered by mapping quality (MAPQ ≥ 30). Peaks were obtained from the “MACS2 callpeak” narrowPeak output files, and overlaps were evaluated strand-agnostically, counting each fragment once if it overlapped any peak. NSC and RSC scores were determined by using the tool: (<https://github.com/kundajelab/phantompeakqualtools>) (2, 3). The scores obtained from all quality metrics were then evaluated based on the recommended threshold values from the ENCODE guidelines (3).

The peaks from the MACS2‐formatted narrowPeak file were filtered based on multiple-testing–corrected significance (q-value < 0.05) and enrichment (signal value ≥ 3.223570, corresponding to the 95th percentile), resulting in 30 significant peaks. Summit positions of each significant peak were then computed as the peak start coordinate plus the peak offset reported by MACS2 callpeak. Additionally, summit positions were stored as 1-bp genomic intervals (GRanges objects) and used for downstream visualization.

The resulting data was then visualized as genome-wide coverage profiles by importing ChIP–seq BedGraph tracks, wherein the coverage scores were normalized to reads per million (RPM) and using the midpoint of each BedGraph interval as the position. The occurrence of the significant peaks was visualized by overlaying the stored summit positions (as described above) as vertical lines on the coverage track.

Locus-level visualization was achieved by first obtaining gene annotations from the GFF3 file of the *C. crescentus* NA1000 genome and then utilizing the gene name if available. If not, the locus tag would be used instead. The data was then subset to a specific region of the genome (NC_011916.1; 3,377,500–3,380,000 bp). In this case, the region corresponds to the highest peak. Summit positions were visualized as vertical lines on the coverage track together with peak intervals as semi-transparent rectangles. Lastly, the genomic context was displayed in the form of a gene track, with separate tracks for genes encoded on the plus and minus strands.

Annotation of peaks from MACS2 was accomplished by mapping them to their closest locus tags of the genome and then performing an additional mapping to an xlxs format file containing all the locus tags of the *C. crescentus* NA1000 genome annotated with COG terms. This was performed for all significant peaks before filtering based on signal value (596 peaks), and after filtering (30 peaks). The “COG genome” file was obtained by submitting the *C. vibrioides* NA1000 genome (NC_011916.1) to “eggNOG-mapper v2” (4, 5), resulting in an xlsx file with all locus tags annotated with Orthologous Groups (OGs) and phylogenies from the EggNOG database (5).

***Motif enrichment analysis***

To identify potential binding motifs, all significant summit positions (30 peaks) were extended by ±100 bp to generate 201 bp genomic windows centered on each summit. Using these genomic windows, sequences were extracted from the *C. vibrioides* NA1000 genome and compiled into a FASTA file and submitted to MEME-ChIP (6) for motif enrichment analysis. All settings for MEME-ChIP were kept at their default values.

***Compositional bias analysis***

Coding DNA sequences (CDSs) were extracted by utilizing the genome (NC_011916.1) and its accompanying annotation file (GFF3) for *C. crescentus* NA1000. To obtain codon usage profiles, non-overlapping frame-dependent triplets were counted. This was done with a relative synonymous codon usage (RSCU) approach. Using a sliding window (step size 1 bp), tetranucleotide frequencies were computed, excluding ambiguous bases, and normalized to the total number of k-mers per CDS. Additionally, frequencies were z-scored by CDS and subjected to principal component analysis (PCA). For each PCA plot, the two PCs that account for the highest variance were visualized, and the centroid of all CDSs was calculated. A 95% confidence ellipse, and highlighted genes of interest were also added to the visualization.

To provide a measure of compositional atypicality potentially indicative of horizontal gene transfer for the genes *higB*, *higA*, and *higX*, we made use of Mahalanobis distance (MD) to quantify the statistical deviation of each CDS and thus the genes of interest from the genomic centroid. Specifically, these two tests were performed: A 2D test based on PC1 and PC2, and a robust k-dimensional test using the Minimum Covariance Determinant (MCD) estimator, wherein the number of PCs is accumulated until a 95% cumulative variance is achieved. Calculation of squared MDs was followed by a comparison against the χ² distribution with degrees of freedom equal to the number of PCs, with outliers defined at the 95% quantile. This was applied to all two tests. Density plots were then used to visualize the genome-wide distributions of MD^2^s with 95% cutoffs, with the genes of interest (*higB*, *higA*, and *higX)* highlighted. To control for multiple hypothesis testing at the genome-wide level, p-values derived from each CDS’s MD2 were adjusted using the Benjamini–Hochberg false discovery rate (FDR) procedure. This was carried out separately for each compositional feature space (RSCU, and tetranucleotide) and for both the 2D (PC1–PC2) and robust k-dimensional tests. Empirical MD cutoffs were then determined by identifying the smallest MD² value among all CDSs with FDR-adjusted q-values ≤ 0.05, thereby defining the genome-wide threshold for compositional atypicality under FDR control.

To quantify the compositional deviation beyond the genome-wide background of the RSCU and tetranucleotide datasets, we utilized squared Mahalanobis distance (MD^2^) values, which had been obtained from the robust MCD analysis. These were then expressed relative to their respective 95% cutoff thresholds and used in making a bar plot of the effect size. For easier interpretability, we opted to use the square root of MD^2^ (√MD^2^), which represents the deviation of each gene from the expected compositional distribution (in a fold change-like manner). Deviation past the 95% robust cutoff would thus be indicated by effect sizes >1. All analyses were performed in R version 4.5.0 (2025-04-11) using RStudio 2024.12.1+563 (“Kousa Dogwood”).

***Phylogenetic and comparative genomic analysis***

*Caulobacterales 16S rRNA phylogenetic tree*

From NCBI, only references and complete genomes belonging to the order Caulobacterales were selected (n=24). 16S rRNA sequences were extracted and aligned using MAFFT in Galaxy, with ”MAFFT flavors” set to ”L-INS-i”. If multiple copies of 16S rRNA were present, only one was extracted to eliminate duplicates. The aligned sequences were subsequently used to create a maximum likelihood (ML) phylogenetic tree using IQ-TREE within Galaxy, with “ultrafast bootstrap replicates” set to 1000. Visualization was then carried out in R with the tree being mid-rooted, bootstrap values (>50) shown at relevant nodes, and organisms colored by genus. The resulting tree would then be used in conjunction with other phylogenetic trees (see below) to assess whether the proteins of the genes identified as outliers also showed atypical conservation patterns.

*16S rRNA tree HigBAX absence/presence matrix*

Genomes were combined into one file and utilized in conjunction with their accompanying GFF files to extract all protein sequences into a single file. Using the tool “NCBI BLAST+ makeblastdb” within Galaxy, a BLAST protein database was made. The database was used for a BLASTx search with three nucleotide query files (*higB*, *higA*, and *higX*). Settings for the BLASTx search were as follows: “Query genetic code” was set to “11. Bacteria and Archaea” and additional columns for the output files were also selected (qlen, slen, saccver, stitle, frames, qcovs, staxids, and sscinames). Lastly, “Maximum hits to consider/show“ was set to 100, and “Maximum number of HSPs” was set to 1.

Tabular output files from the search were filtered by E-value (≤1e-10), percent identity (≥ 35%), and query coverage (≥ 70%). Additionally, rows were discarded if they lacked a resolvable organism key, bit score, E-value, or percent identity. For each gene in each organism, only the single best-scoring BLASTx hit (highest bit score) was kept. If multiple hits shared the same score, one was chosen consistently to avoid duplicates. This produced a nonredundant table of best hits for HigB, HigA, and HigX across all organisms. The data were then used to create a binary matrix based on the presence (1) or absence (0) of each gene per organism. Furthermore, these calls were then summarized by different combination classes and the number of organisms in each of these was counted. The presence/absence matrix was used to create a heatmap and combined with the previous 16S RNA phylogenetic tree.

*Phylogeny of HigBAX*

Query files containing the nucleotide sequence of each gene of interest (*higB*, *higA*, and *higX*) were used for BLASTx searches in Galaxy with most settings left as default except for the following parameters: “Subject database/sequences” was set to “Locally installed BLAST database”, and “Protein BLAST database” was set to “Refseq Protein (13 oct 2024)”. The same settings for “Query genetic code” and additional columns for the output files were kept as stated previously (see previous section).

Hits from the tabular output files were filtered by the same parameters as previously stated (see previous section) and utilized in the creation of a presence/absence (1/0) binary matrix of each gene per organism. Once again, calls were summarized by different combination classes, and the number of organisms in each of these was counted. If a taxid was available, the bacteria were labeled as "<scientific name> [<taxid>]".

FASTA files containing amino acid sequences of three specific combination classes (“HigB only”, “HigA only”, and “HigX only”) were extracted by retrieving the amino-acid sequences of the retained best hits from the NCBI Protein database using their accession numbers. Sequence headers were standardized to the format “scientific name [taxid]”.

Files were submitted to MAFFT within Galaxy followed by IQ-TREE to make a maximum likelihood (ML) phylogenetic tree. ”MAFFT flavors” was set to ”L-INS-i” and IQ-TREE was left at default settings, except for “ultrafast bootstrap replicates” which was set to 1000. Visualization was then carried out in R, with the tree being mid-rooted and bootstrap values (>50) shown at relevant nodes. Organisms were colored by genus or the first section of their name.

***Supplementary Methods references***

1. The Galaxy Community. The Galaxy platform for accessible, reproducible, and collaborative data analyses: 2024 update. Nucleic Acids Research. 2024;52(W1):W83-W94.

**Table S1. HigX peptide abundances, percentage coefficient of variation (CV %) values, abundance ratios (mutants normalized to NA1000 wild type) and p values from quantitative mass spectrometry.**

| Strain | Mean abundances (raw) | CV (%) | Abundance ratios /WT | p value |
| --- | --- | --- | --- | --- |
| Δ*higBA* | 575.3 | 3.83 | 1.126 | 0.259213 |
| Δ*higX* | 285.1 | 5.3 | 0.558 | 0.000134 |
| Δ*lexA* | 733.1 | 9.99 | 1.435 | 0.001817 |
| Δ*lexA* Δ*higBA* | 683 | 12.42 | 1.337 | 0.007723 |
| NA1000 | 510.8 | 11.14 |  |  |

**Table S2. Primers used in this study. Enzyme sites incorporated into primers for cloning purposes are underlined.**

| Primer name | Sequence (5’ – 3’) |
| --- | --- |
| cc3036_wt_nde | AAAACATATGAGCGCTGACGCTTCCAAG |
| cc3036_eco | AAAAGAATTCACTCCGCGTCGTCCGCCA |
| cc3036_noTM_nde | AAACATATGGGCGCCTACTTCCTGCGG |
| cc3036_fusion_eco | AAAGAATTCCTCCGCGTCGTCCGCCAGC |
| 3036_up_bam | AAGGATCCGGAAGCGTCAGCGCTCATGG |
| 3036_up_hind | AAAAGCTTGGTGATCGGTGCTCGATTGG |
| 3036_down_bam | AAGGATCCCTGGCGGACGACGCGGAGTA |
| 3036_down_eco | AAGAATTCTGCCCAGATCGACGCCTTCT |
| higB_pst | AAACTGCAGAACGTCTCCGTCTCGACGA |
| higB_bgl | AAAAGATCTTGTGACATGACGATCAGCGG |
| higX_pst | AAACTGCAGTCAGCGCTCATGGCTCTTG |
| higX_bgl | AAAAGATCTGGGTGACGCCGACTATATCC |
| himup2 | GATATTGCTGAAGAGCTTGGCGGCGAA |
| CC_3036_qrt_fwd | GTTGACGCCAAGGATGGGAT |
| CC_3036_qrt_rev | CATGGTTTGGCCAGTGCCT |
| higB_qrt_fwd | CCGACATGGACCCGCAATTC |
| higB_qrt_rev | GTTCAAGGCTTGGCTAGCGG |
| rpoDfow | GAAGAACTGGCCGAAAAGCT |
| rpoDrev | CGTTCTTGTCCTCGATGAAGTC |

**Table S3. Plasmids used in this study**

| Plasmid name | Description | Reference/construction |
| --- | --- | --- |
| pNPTS138 | colEI ori, M13 ori, oriT, Km^R^, *sacB*, suicide vector for in-frame deletions | M.R.K. Alley (unpublished) |
| pNPTS-Δ*higX* | pNPTS138 derivative to introduce Δ*higX* allele | - Amplification of *higX* upstream region with primers 3036_up_bam and 3036_up_hind, digestion with *Hin*dIII and *Bam*HI - Amplification of *higX* downstream region with primers 3036_down_bam and 3036_down_eco, digestion with *Eco*RI and *Bam*HI - Cloning of the fragments into *Eco*RI and *Hin*dIII sites of pNPTS138 |
| pNPTS-Δ*higBAX* | pNPTS138 derivative to introduce Δ*higBAX* allele | - Digestion of pNPTS-Δ*higBA* (Kirkpatrick *et al,* 2016) with *Eco*RI and *Bam*HI to remove *higBA* downstream region - Amplification of *higX* downstream region with primers 3036_down_bam and 3036_down_eco, digestion with *Eco*RI and *Bam*HI - Cloning of the fragment into pNPTS-Δ*higBA Eco*RI and *Bam*HI sites |
| plac290 | oriV, Tet^R^, lacZ transcriptional fusion vector (low copy) | J. Gober (unpublished) |
| pP*_higBA_*-lac290 | plac290 derivative with promoter of *higBA* inserted upstream of *lacZ* | Kirkpatrick et al, 2016 |
| pJC327 | colEI ori, oriV, Tet^R^, lacZ translational fusion vector | Ardissone et al, 2014 |
| pJC327-P*_higB_-higB::lacZ* | pJC327 derivative with promoter of *higBA* and first 10 codons of *higB* inserted (in-frame) upstream of *lacZ* | - Amplification of *higB* upstream region with primers higB_pst and higB_bgl - Cloning of the fragment into pJC327 *Pst*I and *Bgl*II sites |
| pJC327-P*_higX_-higX::lacZ* | pJC327 derivative with region upstream of *higX* and first 4 codons of *higX* inserted (in-frame) upstream of *lacZ* | - Amplification of *higX* upstream region with primers higX_pst and higX_bgl - Cloning of the fragment into pJC327 *Pst*I and *Bgl*II sites |
| pMT335 | pBBR1 ori, rep, mob, Gent^R^, high copy replicating plasmid for vanillate-inducible gene expression | Thanbichler et al, 2007 |
| pMT335-*higX* | pMT335 derivative, P_van_-*higX* | - Amplification of *higX* coding region with primers cc3036_wt_nde and cc3036_eco - Cloning of the fragment into pMT335 *Nde*I and *Eco*RI sites |
| pMT335-*higX*-GFP | pMT335 derivative, P_van_-*higX-GFP* (HigX-GFP C-terminal translational fusion) | - Amplification of *higX* coding sequence (excluding stop codon) with primers cc3036_wt_nde and cc3036_fusion_eco - Digestion of pMT335-*tipR*-GFP (gift of P. Viollier, unpublished) with *Nde*I and *Eco*RI (removes *tipR* coding sequence) - Cloning of the fragment into *Nde*I and *Eco*RI sites |
| pMT335-*higX*-noTM | pMT335 derivative, P_van_-*higX-noTM* (N-terminal truncation of HigX by replacing amino amino acid Val133 with Met as a new start codon) | - Amplification of partial *higX* coding region with primers cc3036_noTM_nde and cc3036_eco - Cloning of the fragment into pMT335 *Nde*I and *Eco*RI sites |
| pET28a | pBR322 origin, Km^R^, *lacI*, T7-inducible expression vector for N-terminal His6-tagged proteins | Novogen |
| pET28a-higX | pET28a derivative, expresses His6-HigX | - Amplification of *higX* coding region with primers cc3036_wt_nde and cc3036_eco - Cloning of the fragment into pET28a *Nde*I and *Eco*RI sites |
| pET28a-higX-noTM | pET28a derivative, expresses His6-HigX-noTM | - Amplification of partial *higX* coding region with primers cc3036_noTM_nde and cc3036_eco - Cloning of the fragment into pET28a *Nde*I and *Eco*RI sites |
| pHPV414 | oriR6K, Km^R^, himar1 transposon delivery vector | Viollier et al, 2004 |

**Table S4. *Caulobacter crescentus* strains used in this study**

| *C. crescentus* strain | Description/genotype | Reference/construction |
| --- | --- | --- |
| NA1000 | Wild type | Evinger and Agabian, 1997 |
| CLK891 | Δ*higA* | Kirkpatrick *et al*, 2016 |
| CLK113 | Δ*higBA* | Kirkpatrick *et al*, 2016 |
| CLK1659 | Δ*higX* | In-frame deletion of *higX* in NA1000 (retains first six and last six codons of *higX*) |
| CLK1660 | Δ*higBAX* | In-frame deletion of *higBAX* in NA1000 (retains first seven codons of *higB* and last six codons of *higX*, fused in-frame via *Bam*HI site) |
| CLK1203 | Δ*lexA* | Kirkpatrick *et al*, 2016 |
| CLK1204 | Δ*lexA* Δ*higB* | Kirkpatrick *et al*, 2016 |
| CLK1205 | Δ*lexA* Δ*higBA* | Kirkpatrick *et al*, 2016 |
| CLK1574 | Δ*lexA* Δ*higBAX* | In-frame deletion of *higBAX* in CLK1203 |
| CLK1658 | Δ*lexA* Δ*higX* | In-frame deletion of *higX* in CLK1203 |
| CLK133 | NA1000 pP*_higBA_*-lac290 | Kirkpatrick *et al*, 2016 |
| DM217 | Δ*higBA* pP*_higBA_*-lac290 | Kirkpatrick *et al*, 2016 |
| DM223 | Δ*lexA* pP*_higBA_*-lac290 | Kirkpatrick *et al*, 2016 |
| DM224 | Δ*lexA* Δ*higBA* pP*_higBA_*-lac290 | Kirkpatrick *et al*, 2016 |
| CLK1836 | Δ*higBAX* pP*_higBA_*-lac290 | Transformation of CLK1660 with pP*_higBA_*-lac290 |
| CLK1837 | Δ*higX* pP*_higBA_*-lac290 | Transformation of CLK1659 with pP*_higBA_*-lac290 |
| CLK1852 | Δ*lexA* Δ*higBAX* pP*_higBA_*-lac290 | Transformation of CLK1574 with pP*_higBA_*-lac290 |
| CLK1838 | NA1000 pJC327-P*_higB_-higB::lacZ* | Transformation of NA1000 with pJC327-P*_higB_-higB::lacZ* |
| CLK1839 | Δ*higBA* pJC327-P*_higB_-higB::lacZ* | Transformation of CLK113 with pJC327-P*_higB_-higB::lacZ* |
| CLK1840 | Δ*lexA* pJC327-P*_higB_-higB::lacZ* | Transformation of CLK1203 with pJC327-P*_higB_-higB::lacZ* |
| CLK1841 | Δ*lexA* Δ*higBA* pJC327-P*_higB_-higB::lacZ* | Transformation of CLK1205 with pJC327-P*_higB_-higB::lacZ* |
| CLK1842 | Δ*higBAX* pJC327-P*_higB_-higB::lacZ* | Transformation of CLK1660 with pJC327-P*_higB_-higB::lacZ* |
| CLK1843 | Δ*higX* pJC327-P*_higB_-higB::lacZ* | Transformation of CLK1659 with pJC327-P*_higB_-higB::lacZ* |
| CLK1844 | Δ*lexA* Δ*higBAX* pJC327-P*_higB_-higB::lacZ* | Transformation of CLK1574 with pJC327-P*_higB_-higB::lacZ* |
| CLK1853 | NA1000 pJC327-P*_higX_-higX::lacZ* | Transformation of NA1000 with pJC327-P*_higX_-higX::lacZ* |
| CLK1854 | Δ*higBA* pJC327-P*_higX_-higX::lacZ* | Transformation of CLK113 with pJC327-P*_higX_-higX::lacZ* |
| CLK1855 | Δ*lexA* pJC327-P*_higX_-higX::lacZ* | Transformation of CLK1203 with pJC327-P*_higX_-higX::lacZ* |
| CLK1856 | Δ*lexA* Δ*higBA* pJC327-P*_higX_-higX::lacZ* | Transformation of CLK1205 with pJC327-P*_higX_-higX::lacZ* |
| CLK1857 | Δ*higBAX* pJC327-P*_higX_-higX::lacZ* | Transformation of CLK1660 with pJC327-P*_higX_-higX::lacZ* |
| CLK1858 | Δ*higX* pJC327-P*_higX_-higX::lacZ* | Transformation of CLK1659 with pJC327-P*_higX_-higX::lacZ* |
| CLK1859 | Δ*lexA* Δ*higBAX* pJC327-P*_higX_-higX::lacZ* | Transformation of CLK1574 with pJC327-P*_higX_-higX::lacZ* |
| CLK1834 | NA1000 pMT335-*higX*-GFP | Transformation of NA1000 with pMT335-*higX*-GFP |
| CLK1835 | Δ*lexA* Δ*higX* pMT335-*higX*-GFP | Transformation of CLK1658 with pMT335-*higX*-GFP |
| CLK196 | NA1000 pP*_higBA_*-lac290 pMT335 | Kirkpatrick *et al*, 2016 |
| CLK1508 | NA1000 pP*_higBA_*-lac290 pMT335-*higX* | Transformation of CLK133 with pMT335-*higX* |
| CLK1661 | NA1000 pP*_higBA_*-lac290 pMT335-*higX*-noTM | Transformation of CLK133 with pMT335-*higX*-noTM |
| CLK234 | Δ*higBA* pP*_higBA_*-lac290 pMT335 | Kirkpatrick *et al*, 2016 |
| CLK1510 | Δ*higBA* pP*_higBA_*-lac290 pMT335-*higX* | Transformation of DM217 with pMT335-*higX* |
| CLK1662 | Δ*higBA* pP*_higBA_*-lac290 pMT335-*higX*-noTM | Transformation of DM217 with pMT335-*higX*-noTM |
| CLK1514 | Δ*lexA* pP*_higBA_*-lac290 pMT335 | Transformation of DM223 with pMT335 |
| CLK1512 | Δ*lexA* pP*_higBA_*-lac290 pMT335-*higX* | Transformation of DM223 with pMT335-*higX* |
| CLK131 | NA1000 pMT335 | Kirkpatrick et al, 2016 |
| CLK1813 | Δ*lexA* pMT335 | Transformation of CLK1203 with pMT335 |
| CLK1663 | Δ*lexA* Δ*higB* pMT335 | Transformation of CLK1204 with pMT335 |
| CLK1664 | NA1000 pMT335-*higX* | Transformation of NA1000 with pMT335-*higX* |
| CLK1665 | Δ*lexA* pMT335-*higX* | Transformation of CLK1203 with pMT335-*higX* |
| CLK1666 | Δ*lexA* Δ*higB* pMT335-*higX* | Transformation of CLK1204 with pMT335-*higX* |
| CLK1790 | Δ*lexA* Δ*higBA* CCNA_03248::*himar1* | *himar1* insertion into coding sequence of CCNA_03248, transduced with ΦCr30 into CLK1205 |
| CLK1791 | Δ*lexA* Δ*higBA* CCNA_00669::*himar1* | *himar1* insertion into coding sequence of CCNA_00669, transduced with ΦCr30 into CLK1205 |
| CLK1792 | Δ*lexA* Δ*higBA* P*_amiC_*::*himar1* | *himar1* insertion upstream of *amiC* coding sequence, transduced with ΦCr30 into CLK1205 |
