## Supplementary figures and images for "An ancestral transmembrane transcription factor couples cell envelope regulation and the SOS response in *Caulobacter crescentus*"

### Supplemental Figures

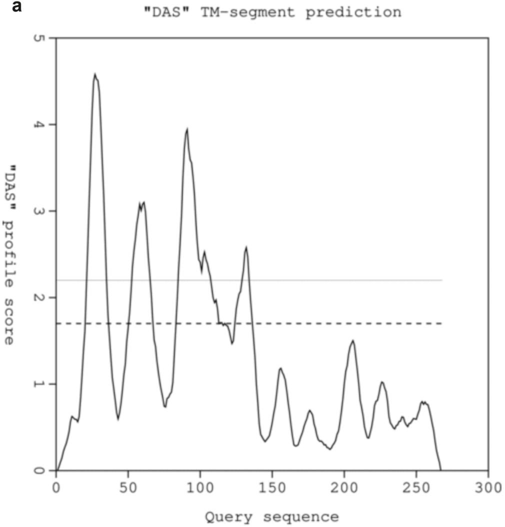

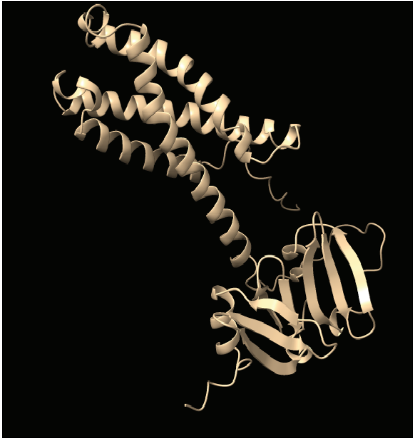

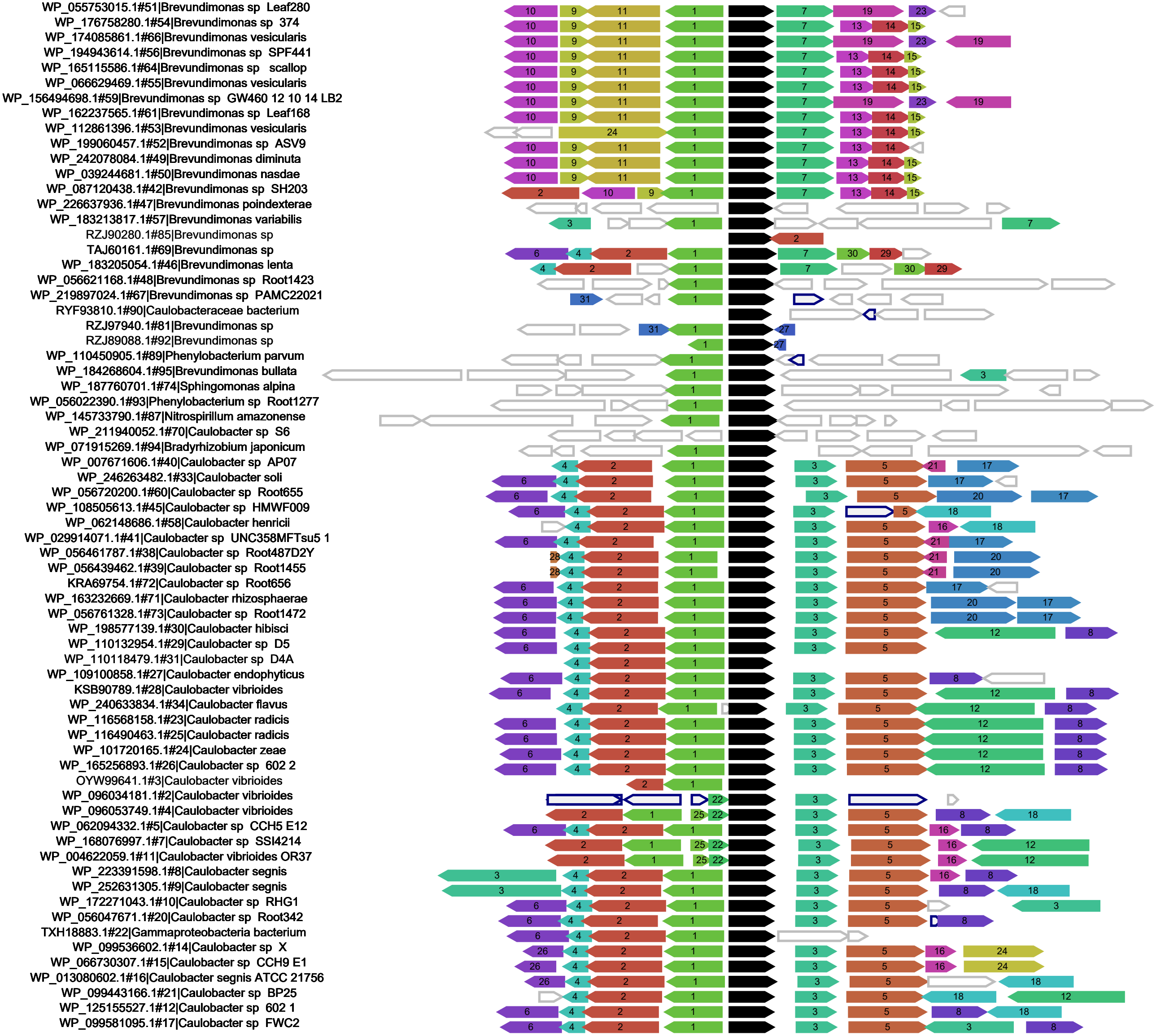

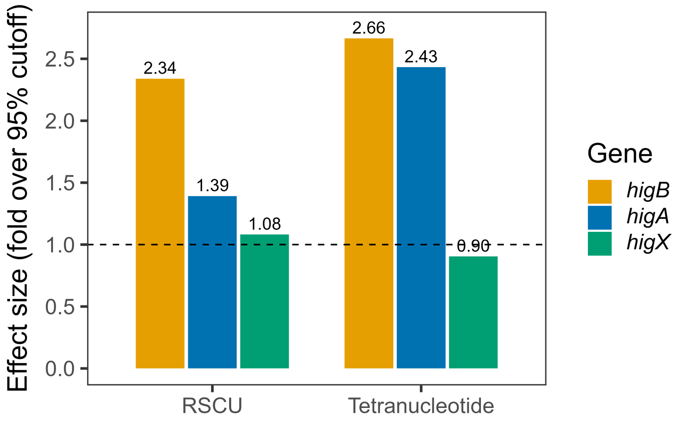

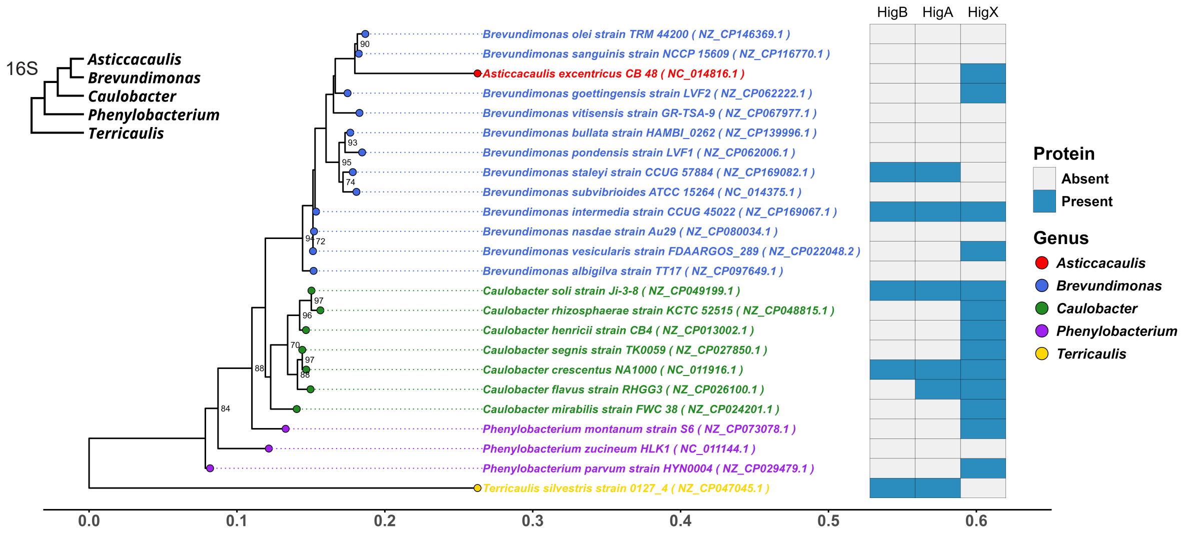

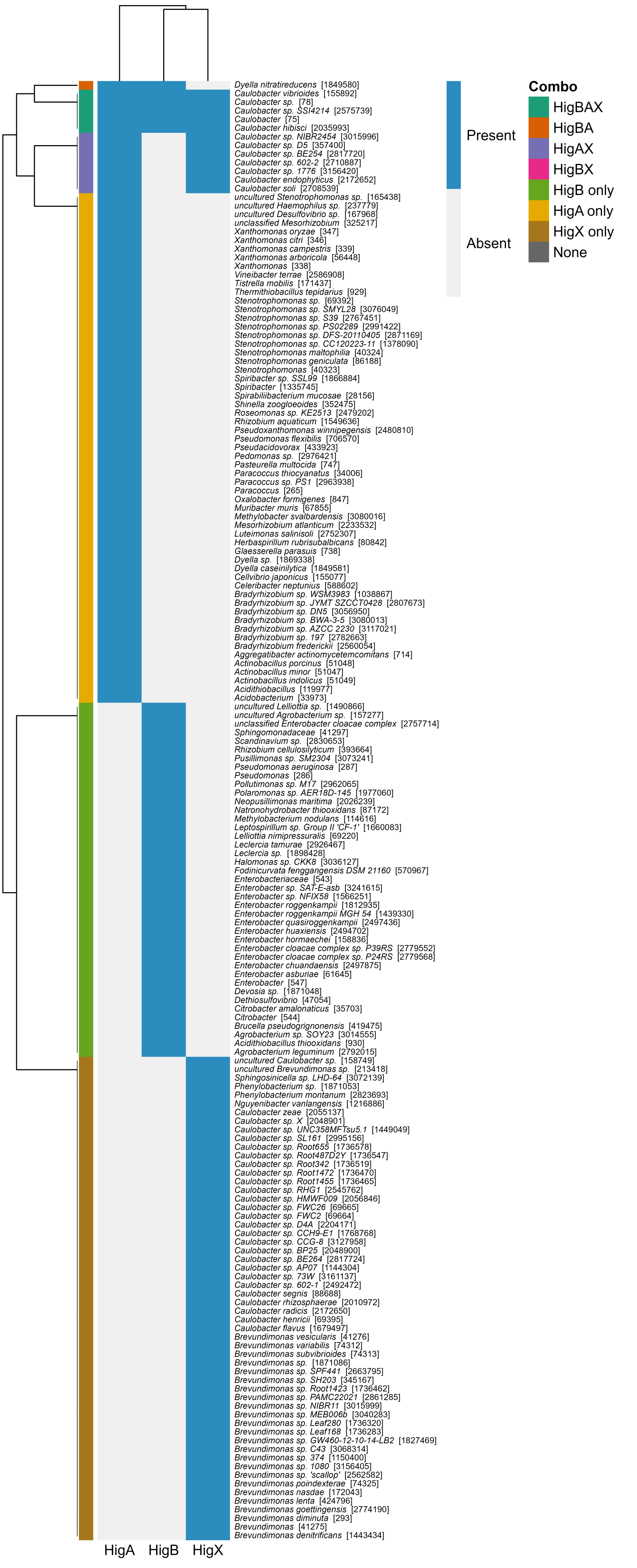

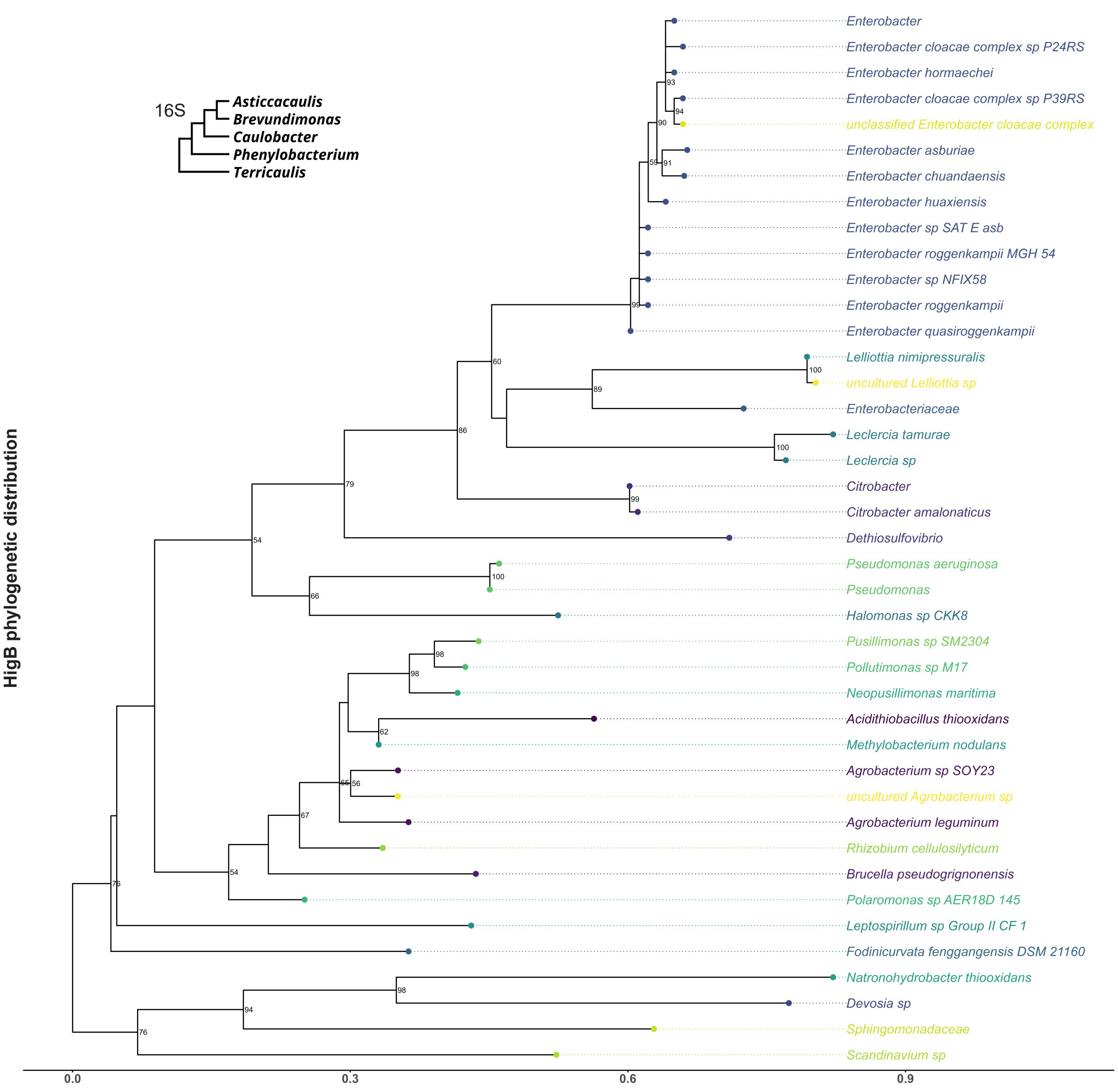

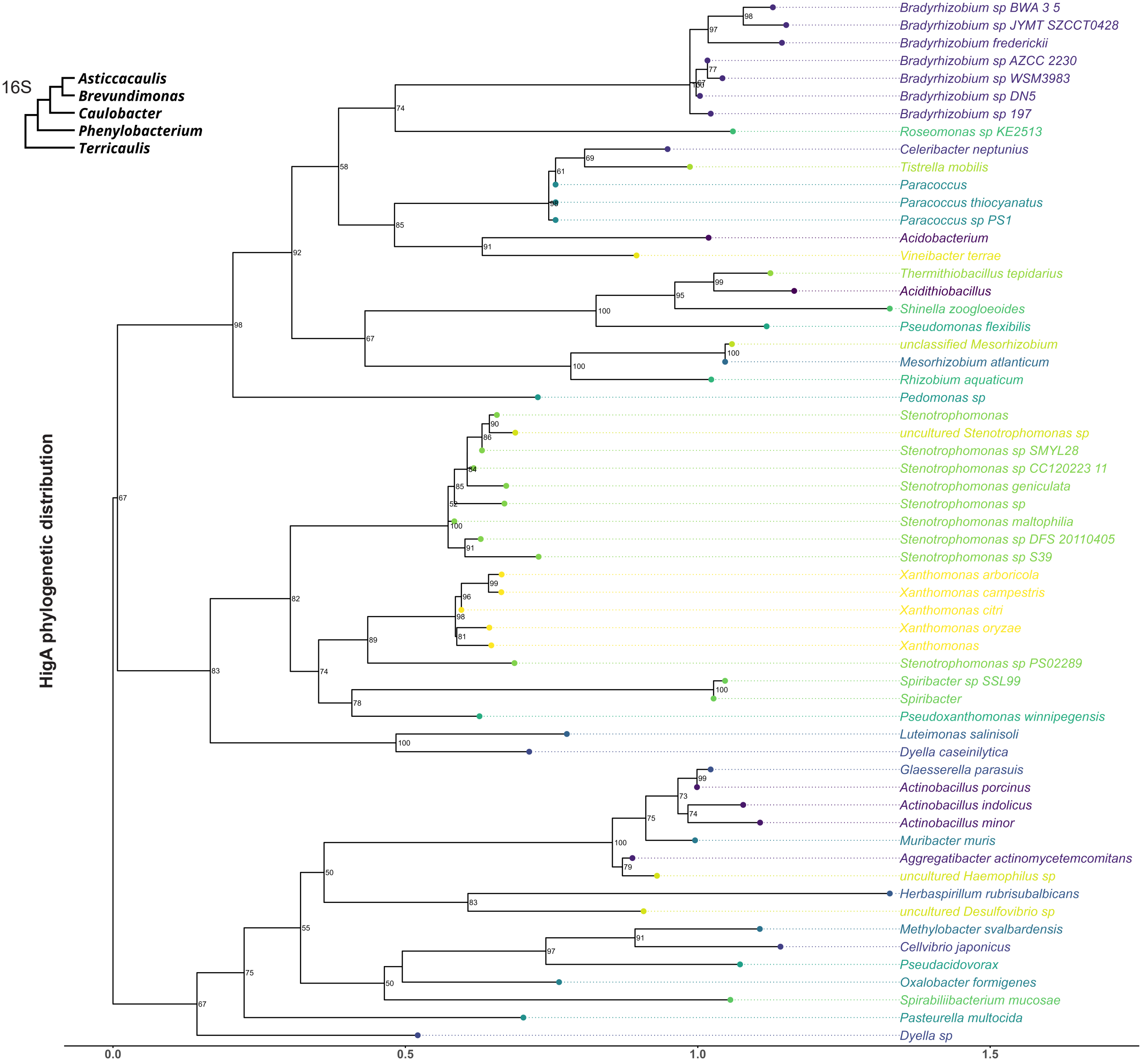

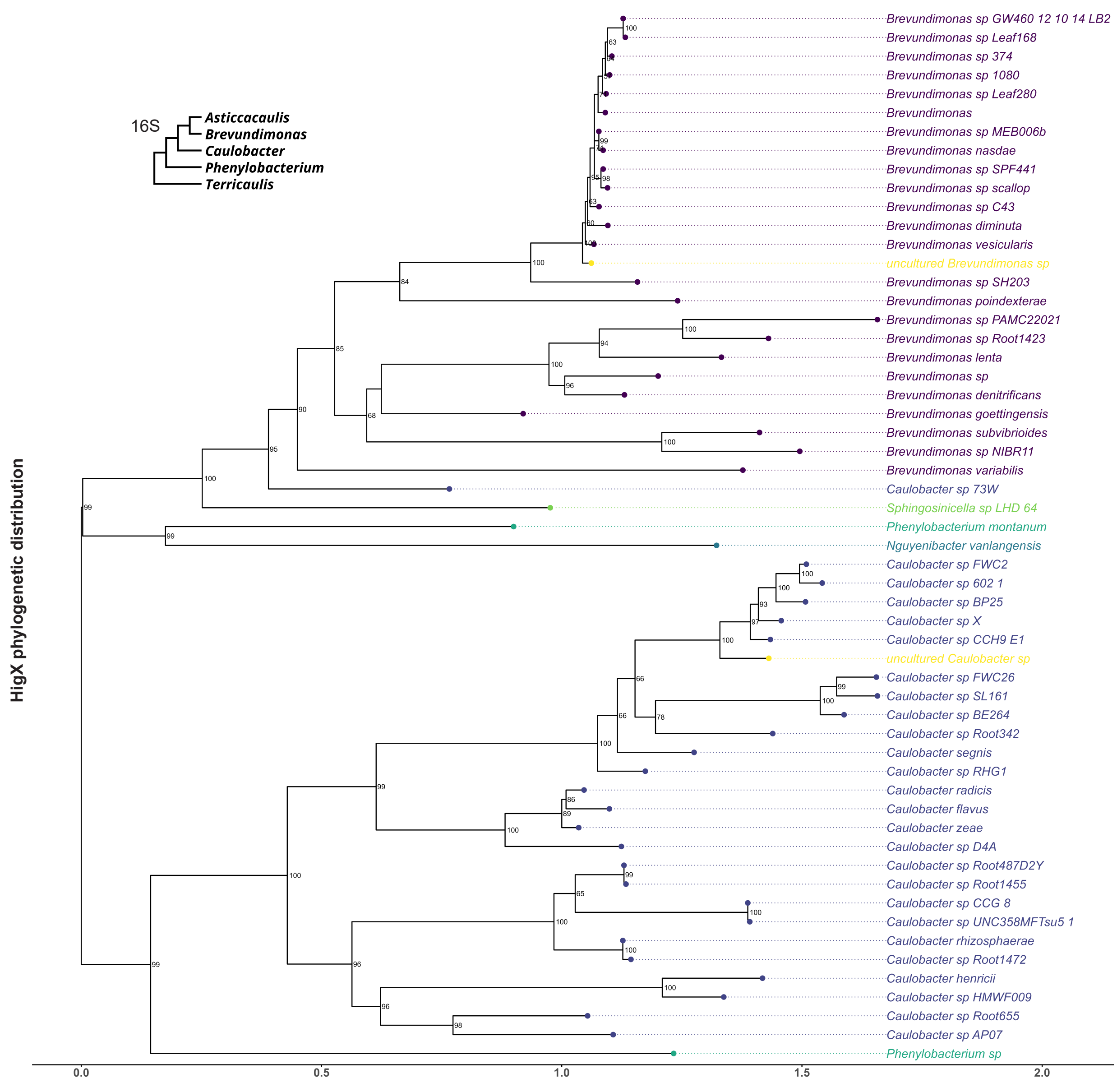
